# Functional definition of the *Drosophila* airway progenitor field through overlapping compensatory regulators

**DOI:** 10.64898/2026.03.18.712720

**Authors:** Ryo Matsuda, Chie Hosono, Kaoru Saigo, Christos Samakovlis

**Affiliations:** SciLifeLab and Department of Molecular Biosciences, The Wenner-Gren Institute, Stockholm University, 10691, Stockholm, Sweden; Department of Biophysics and Biochemistry, School of Science, the University of Tokyo, Tokyo, Japan; Cardiopulmonary Institute (CPI), Member of the German Center for Lung Research (DZL), Justus Liebig University, Giessen, Germany

## Abstract

Tubular organs present a common solution to fluid transport in multicellular organisms. They often arise by an initial bulging of flat epithelial progenitor cells, which then undergo branching morphogenesis. Here, we present 3 cooperative programs fully defining the progenitor field of the *Drosophila* airways (the tracheal system) and their roles in early morphogenesis linking the radial pattern of the 2-dimensional (2D) field to the proximo-distally patterning of the 3D tubes. We previously showed that extrinsic Hedgehog (Hh) and intrinsic POU-Homeobox TF Ventral-veinless(Vvl)/Drifter/U-turn drive the transcriptional program toward the distal airway cell identity at the expense of a proximal program specified by the GATA TF *grain* (*grn*). Both programs require the basic-HLH-POU TF *trachealess* (*trh*) (Matsuda et. al, 2015) but *trh* is not essential for primordia invagination. Now, we report the genetic requirements of airway progenitor invagination. We show that in *hh vvl* double mutants, the oval-shaped primordia frequently remain at the 2D plane, retaining *trh* expression in a *grn* dependent manner. Therefore, *hh* and *vvl* are the principal regulators of progenitor invagination independent of *trh*. Each of the 3 transcriptional regulators, Trh, Vvl and Grn fulfill only complementary or compensatory functions in transcription and morphogenesis, but their combinations functionally define the airway progenitor field. We further provide a comprehensive description for allocation of the airway progenitors on the body coordinates, involving dorsal Decapentaplegic/BMP signaling along the dorso-ventral axis and subsequent radial EGFR signaling along the proximo-distal axis. The presence of 3 complementary, regulatory programs in early gene expression and morphogenesis of the simple *Drosophila* airways may reflect the vital needs for respiration, and their influence on the evolution of various strategies in tubular organ development.

## Introduction

The reiterative tube formation and ramification in airways and lungs generate the pulmonary-vascular network to efficiently supply oxygen to the whole body. The airway tubes allow the airflow to alveolar structures, where blood cells inside the fine vascular tubes exchange carbon dioxide with oxygen to deliver it to the body (Herriges and Morrisey 2014; Potente and Makinen 2017; Kishimoto and Morimoto 2021). Surprisingly, a similar pulmonary-blood cell connection is also found in the fruit fly *Drosophila*. There, reiterative ramification of the epithelial tubes (the tracheal system) allows the airflow inside the body that directly delivers oxygen to the target cells (Manning and Krasnow 1993; Samakovlis et al. 1996a; Hu and Castelli-Gair 1999; Ghabrial et al. 2003), while the blood cells in the open circulatory system transiently attach to the fine airways to boost oxygen transport (Shin et al. 2024).

The lungs and the *Drosophila* airways both derive from 2D- primordial cell fields (Romero et al. 2025). Lung formation initiates as tube evaginations from the foregut epithelium (Herriges and Morrisey 2014; Kishimoto and Morimoto 2021). Distal ramification of these buds and further differentiation into distal airways and alveoli involve FGFR activation by extrinsic FGFs (Min et al. 1998; Weinstein et al. 1998; Arman et al. 1999; Sekine et al. 1999; Brownfield et al. 2022; Jones et al. 2022; Sountoulidis et al. 2023; Sountoulidis et al. 2026). Similarly, *Drosophila* airway formation initiates from cell primordia specified on the ectodermal plane, each invaginating to form a primitive cavity (Perrimon et al. 1991; Manning and Krasnow 1993; Samakovlis et al. 1996a; Hu and Castelli-Gair 1999). Extrinsic FGF/Branchless (Bnl) activates FGFR/Breathless (Btl) to guide branch ramification and network connection (Klambt et al. 1992; Guillemin et al. 1996; Samakovlis et al. 1996a; Samakovlis et al. 1996b; Sutherland et al. 1996) (Figure 1A). Accordingly, highly ramified fine tubes are generated around each branch terminus that serve for oxygen transport.

**Figure 1.**
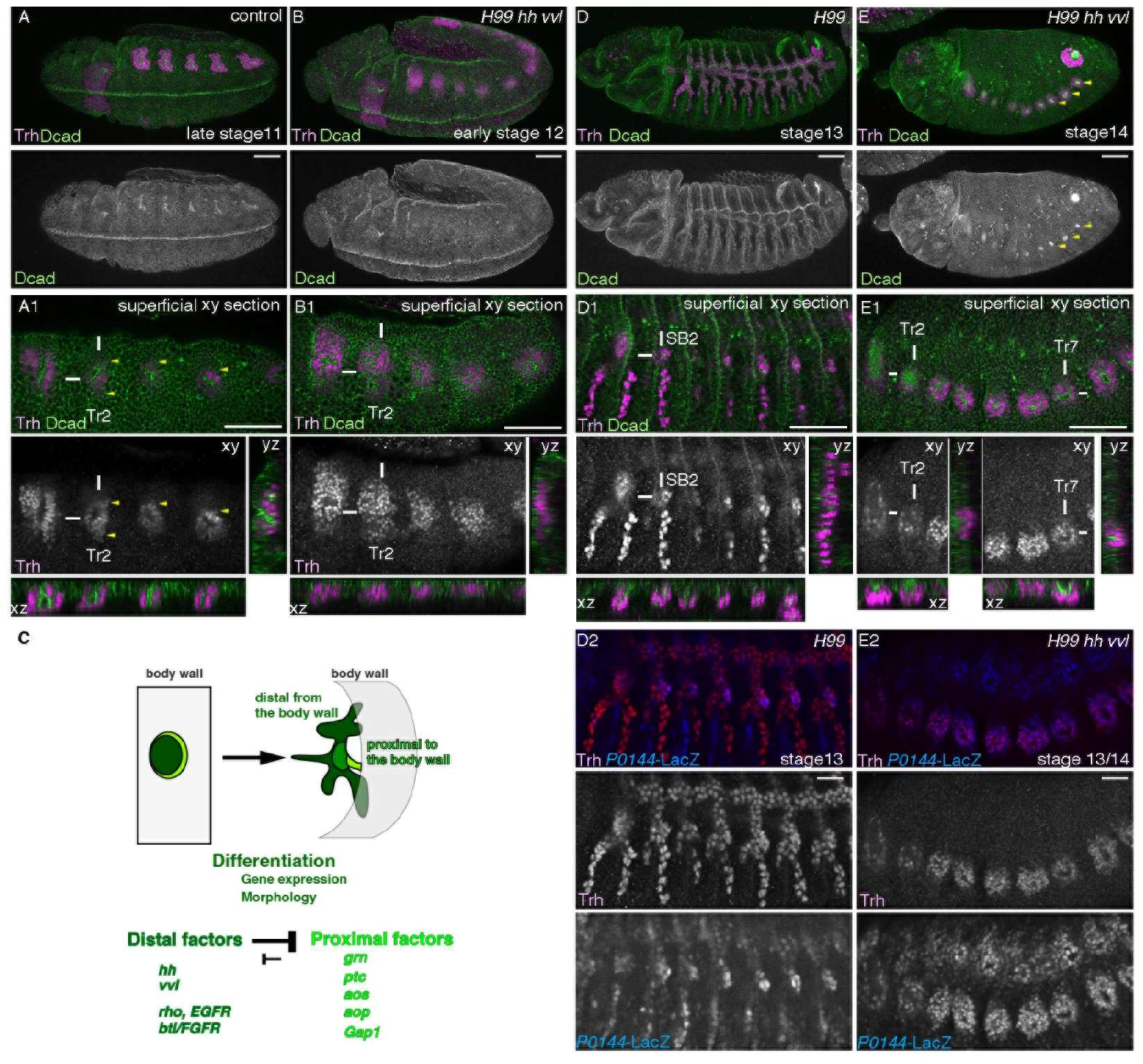
*hh* and *vvl* empower invagination of the airway progenitors. (A-B, D-E) Progenitor invagination and Trh expression in the control (A, D) or *H99 hh vvl* mutants (*H99 hh^13C^ vvl^utH599^* homozygotes) (B, E), visualized by staining for D-cadherin/Dcad (Shotgun) (Oda et al. 1994; Tepass et al. 1996). A, B, D, D2 and E2 are projections while A1, B1, D1, E1 represent xy- yz- and xz- sections of embryos shown in A, B, D and E, respectively. White lines mark the places of yz- and xz-sectioning. Signals are enhanced in sectional views. In the control (A, A1), progenitor invagination and tubulogenesis are evident as early as late stage 11. In *H99 hh vvl* mutants at stage 12 (B, B1), apical enrichment of Dcad signals and apical extrusion of Trh^+^ nucleus are irregularly detected. However, persistent progenitor invagination to form deep cavities rarely happens. Note that in A1, Trh is detected in the epidermal layer (yellow arrowheads) but our reagents do not allow accurate discrimination between epidermal or airway progenitors. At stages 13/14, compared to the control (D, D1, D2), in *H99 hh vvl* mutants (E, E1, E2), apical enrichment of Dcad signals and apical constriction are persistently detected around the central parts of each primordium, that eventually form shallow cavities (yellow arrowheads in E). Trh and strong *P0144*-LacZ signals overlap mainly in the spiracular branch in the control (D2) whereas in *H99 hh vvl* mutants (E2), *P0144*-LacZ^+^ progenitors occupy the whole progenitor areas. Trh is detected in smaller areas than *P0144*-LacZ, probably due to the gradual loss of Trh expression in the periphery. Note that in A1, B1, D1 or E1, the numbers of tracheal metamer (Tr) or spiracular branch (SB) are labelled. Scale bar is 50 μm (C) A schema showing that invagination transforms the centro-peripheral patterning of the primordium into the proximo-distal patterning of the tubes. Extrinsic Hh and intrinsic Vvl promote the distal gene expression program (dark green domain) whereas *rho* and *btl/FGFR* activating morphogenesis and distal gene expression programs. The proximal factors including *grn* promote the proximal gene expression program (light green domain).

*trh* is the earliest TF-encoding gene marking the *Drosophila* airway primordia and the mature airways (Isaac and Andrew 1996; Wilk et al. 1996). *trh* has been regarded as the master TF of the *Drosophila* airways because tubes as well as differentiation markers are not detected in *trh* mutants at later stages (Isaac and Andrew 1996; Wilk et al. 1996; Brodu and Casanova 2006; Sotillos et al. 2010; Chung et al. 2011; Matsuda et al. 2015b). However, in *trh* null mutants, expression of *rhomboid* (*rho*) (Bier et al. 1990), a protease activating the EGFR ligand Spitz (Schweitzer et al. 1995) initiates in the primordia (Ogura et al. 2018) but its propagation (Matsuda et al. 2015b) fails (Ogura et al. 2018). Consistent with the role of *rho* in promoting primordial invagination (Brodu and Casanova 2006; Nishimura et al. 2007), *trh* mutants initiate primordia invagination but neither sustained invagination nor maintained tube structures (Kondo and Hayashi 2019). The initiation of invagination and the early localized *rho* expression in *trh* mutants suggest additional regulators, other than *trh* that are critically responsible for gene expression in airway primordia and for tubulogenesis.

We had previously shown that the proximo-distal axis of the *Drosophila* airway tubes is generated from the centro-peripheral patterning of the 2D primordia (Matsuda et al. 2015b). One set of genes, the distalizing factors, *hh*, *vvl*, *rho* and *btl/FGFR* cooperate to realize the distal gene expression program at the expense of the proximal one, whereas another set, the proximal factors including *grn*, negative regulators of *EGFR* signaling and *hh* signaling realize the proximal gene expression program (Figure 1C). The two programs establish distinct proximal and distal cellular domains in the branching network and maintain *trh* expression (Matsuda et al. 2015b). Two of the distalizing factors, *rho* and *btl/FGFR* promote primordia invagination but are not essential for tubulogenesis (Brodu and Casanova 2006; Matsuda et al. 2015b), indicating that essential regulators of primordial morphogenesis have been elusive.

Here, we report that in the absence of the remaining two distalizing factors *hh* and *vvl*, the trachea primordia cells frequently fail invagination and are only detected on the 2D plane. Therefore, extrinsic Hh and intrinsic Vvl represent the missing factors that cooperatively promote primordia invagination. Although *trh*, *vvl* or *grn* alone cannot define the airway progenitors, we propose that combinations of the three TFs can intrinsically define the airway progenitors, considering their subsequent roles on gene expression and morphogenesis. Even in the simple *Drosophila* airways, combinations of multiple factors induce the organ progenitor field and subsequent tubulogenesis to cope with physiological stress of respiration in a terrestrial environment.

## Results and Discussion

### The distalizing factors *hh* and *vvl* drive airway primordia invagination independent of *trh*

Among the distalizing factors, *hh*, *vvl*, *rho* and *btl/FGFR* (Figure 1C), *hh vvl* double mutants show more extensive distal-to-proximal gene expression conversion than the loss of both *rho* and *bnl/FGF-btl/FGFR* signaling (Matsuda et al. 2015b). As *EGFR* (also known as *torpedo/top* or *faint little ball)* and *btl/FGFR* signaling also orchestrate the distal morphogenetic processes, including primordial cell invagination (Llimargas and Casanova 1999; Brodu and Casanova 2006; Nishimura et al. 2007; Kondo and Hayashi 2013) and subsequent branching (Glazer and Shilo 1991; Klambt et al. 1992; Sutherland et al. 1996; Llimargas and Casanova 1999; Matsuda et al. 2015b), *hh* and *vvl* are expected to control the distal morphogenetic program as well. In the absence of *hh* and *vvl*, *rho* expression is detected at early stages sporadically and weakly (Figure 1-figure supplement 1A, B) and *btl* expression initiates in the primordia (Matsuda et al. 2015b), but both fade away shortly thereafter (Matsuda et al. 2015b). Despite the initial expression of *rho* and *btl/FGFR* in *hh vvl* mutants, *hh vvl* embryos show more severe branching phenotypes than the *rho btl* mutants (Matsuda et al. 2015b).

We thus investigated potential invagination defects of *trh* positive cells in *hh vvl* double mutants. We monitored *trh* expression by enhancer trap insertion (*trh*-LacZ) and *trh* transcripts by *in situ* hybridization (Figure 1-figure supplement 1C). Perdurance of LacZ/beta-Galactosidase protein allows tracing the cells that started *trh* transcription even after its transcription ceases. In *hh vvl* double mutants, we detected aberrant expression of both *trh* RNA and *trh*-LacZ persisting on the epidermal plane at stage 12 (Figure 1-figure supplement 1D, E). However, massive ectodermal apoptosis precluded analysis at later stages. Therefore, we switched to *H99 hh vvl* triple mutants, where apoptosis is suppressed by deletion of major pro-apoptotic genes (White et al. 1994).

Compared to the control where primary branching after progenitor invagination is evident by late stage 11, we often detected failure in invagination in the mutants (Figure 1A, A1, B, B1). Out of 54 tracheal metameres (Tr1-9 of 6 embryos between late stage 11 and early stage 12), a small part of the progenitors invaginated to form tubes in only 4 metameres. We also sporadically detected apical constriction and apical extrusion of nuclei, in addition to indentations of a group of progenitors in 4 metameres. In contrast, by stage 15, small cavities were detected in most metameres of *H99 hh vvl* mutants. However, Trh+ cells did not invaginate completely and there were Trh+ cells still sitting in 2D planar configuration around the cavities. All these cells were positive for the proximal marker *P0144*-LacZ (Figure 1D1, D2, E1, E2), suggesting preceding morphogenetic activities. Indeed, 35 out of 81 tracheal metameres in late stage 13 and stage 14 embryos (n=9), had formed tubes, whereas the remaining 36 showed partial indentations with progenitor cells simultaneously exhibiting apical constriction and apical extrusion of nuclei. The partial invagination defects in *H99 hh vvl* mutants at early stages suggest that there are mechanisms other than *hh* and *vvl* that promote tubulogenesis. Progenitor mitosis or residual expression of *rho* or *<u>btl/FGFR</u>* in *H99 hh vvl* mutants may account for this (Llimargas and Casanova 1999; Brodu and Casanova 2006; Nishimura et al. 2007; Kondo and Hayashi 2013). Moreover, the late invagination of the *H99 hh vvl* mutant progenitors suggests that the proximal cell types can become internalized by a mechanism independent of distal cell type differentiation. Because in wild type, the spiracular branches elongate significantly after invagination (Figure 1-figure supplement 1F, G), we speculate that the internalization of the proximalized progenitors in *H99 hh vvl* mutants may be promoted by this unexplored SB elongation mechanisms.

**Figure 2.**
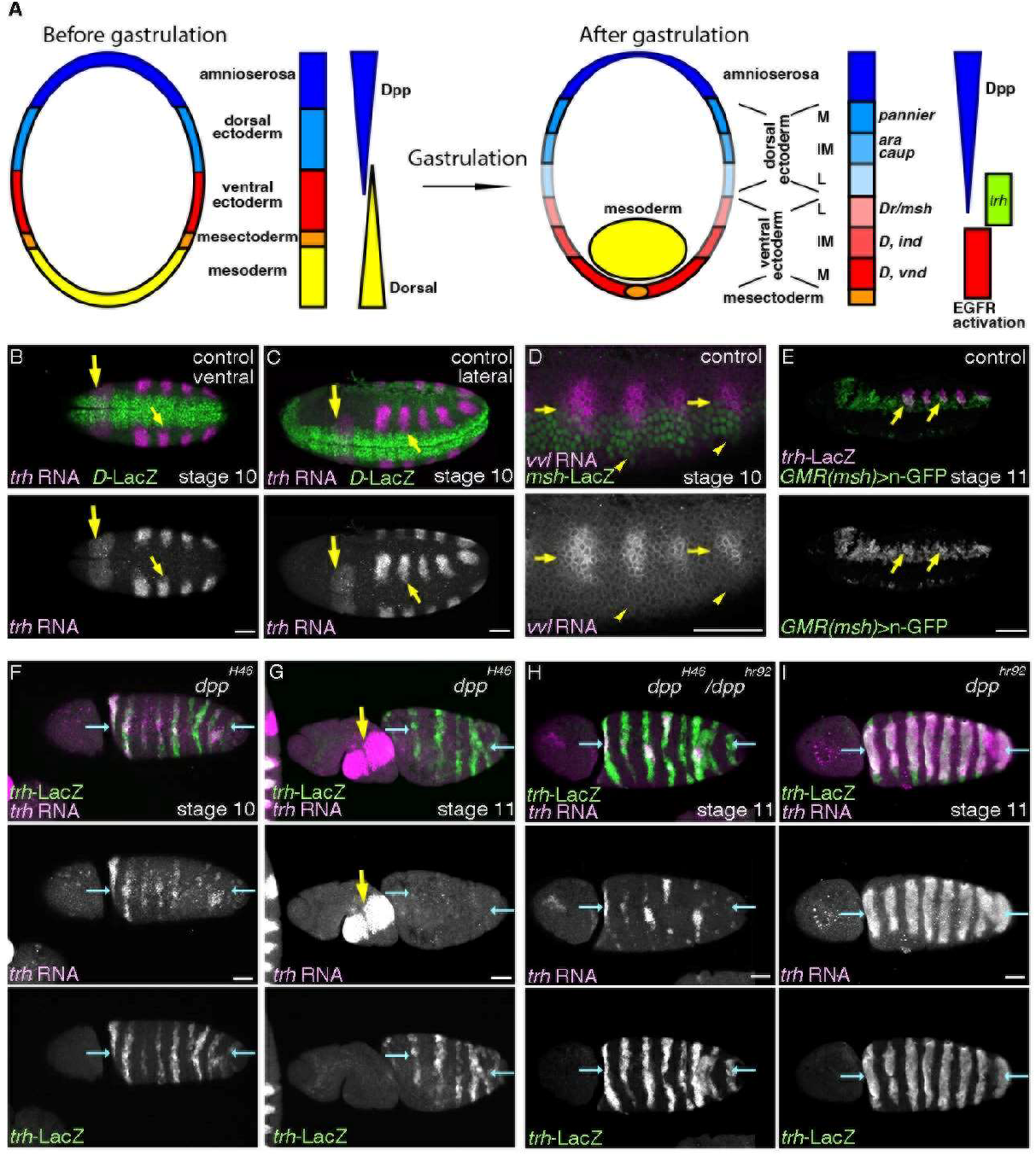
Intermediate Dpp/BMP activity promotes *trh* expression and the airway progenitors. (A) A schema showing that the embryonic trunk is divided into discrete domains along the D-V axis. Out of the 5 domains (amnioserosa, dorsal ectoderm, ventral ectoderm, mesectoderm and mesoderm), mesoderm invaginates during gastrulation. Graded activities of a TF *dorsal* orchestrates the ventral domains whereas Dpp/BMP signaling directs the dorsal domains. Differential expression of TFs subdivides both the dorsal and the ventral ectoderm into medial (M), intermediate (IM) and lateral (L) columns. EGFR is dynamically activated to establish the ventro-intermediate column (Yagi et al. 1998). Expression of TFs marks the column of dorso-medial (*pannier*), dorso-intermediate (*araucan/ara, caupolican/caup*), ventrol-lateral (*Dr/msh*), ventro-intermediate (*Dicheaete/D, intermediate neuroblasts defective/ind*), ventro-medial (*D, ventral nervous system defective/vnd*). (B-E) Distribution of *trh* and *vvl* transcripts relative to the ectodermal subdivision along the D-V axis. (B-C) Ventral and lateral views of stage 10 embryos. The ventral limit of *trh* expression abuts the dorsal limit of *D*-LacZ expression (small yellow arrows) which marks the ventro-medial and the ventro-intermediate columns. Large arrows show *trh* expression in the salivary gland primordia. (D-E) *vvl* expression at stage 10 straddles the border between the dorsal ectoderm and the *Dr/msh*-LacZ positive ventro-lateral ectoderm (D, yellow arrows) whereas the ventral limit of *vvl* expression is some cells away from the ventral limit of *Dr/msh*-LacZ expression (D, yellow arrowheads). An enhancer fragment of *Dr/msh* active in the ventro-lateral column marks the ventral parts of the airway tubes at stage 11 (E, yellow arrows). (F-I) Expression of *trh*-LacZ and *trh* transcripts in allelic series of *dpp* mutants. Dorsal views of embryos where the presumptive dorsal midline are marked with blue arrows. In *dpp* null mutants (F, G), *trh* expression detected with *trh*-LacZ or *trh* RNA expands to the dorsal midline (blue arrows). However at later stages, *trh* RNA is not maintained though *trh*-LacZ positive cells remain (G). Note that in *dpp* null mutants the body is twisted so that *trh* expression in the salivary gland primordia (yellow arrowheads) and in the airway primordia are simultaneously visible. In the milder condition (H), *trh* RNA is sporadically maintained near the dorsal midline whereas in *dpp* hypomorph homozygotes (I), *trh* RNA is detected in many of the expanded progenitor areas. Scale bar is 50 μm

Taken together, the two distalizing factors *hh* and *vvl* define the central/distal cell identity, controlling both gene expression and cell invagination. It follows that 3 intrinsic TFs, Trh, Vvl and Grn cover the major functional aspects of airway progenitor differentiation, transcriptionally and morphologically.

### *dpp/BMP* specifies the airway progenitors along the D-V axis

We further investigated upstream determinants of the primordial field defined by the expression of these 3 TFs. We first allocated their expression domains along the D-V axis of the embryo. By mid-embryogenesis (stage 10-12), the surface of the embryonic trunk becomes largely divided into 3 sectors along the D-V axis, amnioserosa, the dorsal and the ventral ectoderm (Figure 2A) (Wharton et al. 1993; von Ohlen and Doe 2000). Both the dorsal and the ventral ectoderm are further subdivided into 3 parts along the D-V axis, medial, intermediate and lateral (Figure 2A) (von Ohlen and Doe 2000). The dorsal intermediate column marked with *caupolican (caup)/araucan (ara)* spans several cells located dorsally to the *trh*-LacZ positive spiracular branches (Figure 2-figure supplement 1A), whereas the Grn-GFP expression domain, which also marks the most proximal cells of spiracular branches (Figure 2-figure supplement 1C, D) extends ventrally from the dorso-lateral ectoderm (Figure 2-figure supplement 1B). On the other hand, the ventral limits of the initial *trh* expression at stage 10 abut *Dichaete*-LacZ (*D*-LacZ) expressing cells in the intermediate and medial columns of the ventral ectoderm (Figure 2B-C) (Zhao and Skeath 2002). Thus, the initial *trh* expression straddles from the ventro-lateral column to halfway to the dorso-lateral column. Compared to *trh* expression, *vvl* expression in the airway primordia is more restricted (Figure 2D, Figure 2-figure supplement 1E, F). *vvl* expression straddles the boundary of the dorsal and the ventral ectoderm demarcated by *Drop (Dr)/muscle homeobox (msh),* whereas the ventral limit of *vvl* expression is far from the ventro-intermediate column (Figure 2D) (von Ohlen and Doe 2000). Consistently, the ventro-lateral ectoderm enhancer of *Dr/msh* (Pfeiffer et al. 2008; Manning et al. 2012) marks the ventral parts of the invaginated airway progenitors (Figure 2E).

In the absence of Dpp/BMP, the dorsal 2 sectors, amnioserosa and the dorsal ectoderm take the cell differentiation program of the ventro-lateral ectoderm (von Ohlen and Doe 2000). Previous work showed that the intermediate level of Dpp/BMP signaling is responsible for establishing the expression patterns of *vvl* and *grn* for early airway development (Matsuda et al. 2015b). Because Dpp/BMP signaling is a major determinant of the dorsal-ventral embryonic axis, we focused on its role on setting the *trh* expression domain. The initial *trh* expression straddles the ventro-lateral ectoderm in control embryos (Figure 2B, C) but expands to the dorsal midline in *dpp* null mutants (Figure 2F) (Isaac and Andrew 1996). This expression is extinguished by stage 12 in *dpp* null mutants (Figure 2G), consistent with the role of intermediate Dpp/BMP signaling on the expression of *vvl* and *grn*, which in turn maintain *trh* expression (Matsuda et al. 2015b). At milder inactivation conditions of *dpp/BMP* hypomorphic mutants (*dpp^H46^/dpp^hr92^*trans-heterozygotes) (Wharton et al. 1993), *trh* maintenance, if any, occurs only around the dorsal midline, where reduced Dpp/BMP activity levels are presumed to be sporadically present (Figure 2H). In even milder conditions of *dpp/BMP* inactivation in *dpp^hr92^* hypomorphs (Wharton et al. 1993), *trh* maintenance occurs in the more ventral cells as well (Figure 2I). However, we note that concomitant with reduction of Dpp/BMP activities, more cells in the ventral part of the initial trh expression domain fail to maintain *trh* expression (Figure 2F-I, Figure 2-figure supplement 1G, H). Loss of the ventral *trh* expression appears to occur also in wild type embryos (Figure 2-figure supplement 1G), possibly reflecting that the most ventral cells are farthest from the Dpp/BMP source.

The distal and the proximal progenitors are specified in *CycA* mutants (Matsuda et al. 2015b), where the last progenitor mitosis does not occur (Beitel and Krasnow 2000). As the distal cells of the branched airway tree are double positive for *trh* and *<u>vvl</u>*(Matsuda et al. 2015b) and occupy 90% of the mature airways (Beitel and Krasnow 2000), we speculate that Trh^+^ primordial cells within, or bordering the Vvl^+^ expression domain are the airway progenitors straddling the canonical dorsal-ventral ectoderm boundary. The remaining Trh^+^ cells would be epidermal and fail to invaginate. We conclude that optimal intermediate Dpp/BMP activities below the repressive level of action specify the airway progenitors along the D-V axis.

### The radial EGFR signaling primes the airway progenitors and realizes airway differentiation along the P-D axis

Airway cells expressing late differentiation markers are reduced in number in *H99 EGFR btl/FGFR* mutants, where anti-apoptotic functions of *btl/FGFR* and *EGFR* are compensated by *H99* deficiency (Matsuda et al. 2015b). These missing cells prompted us to investigate the potential roles of the two RTKs on earlier *trh* expression.

*trh* expression is comparable in the distal and the proximal cells at stage 12 in *H99 btl/FGFR* mutants (Figure 3A). In contrast, *trh* expression in the proximal area becomes reduced or extinguished at stage 12 in *H99 EGFR* mutants (Figure 3B). Moreover, *trh* expression in the main airways is very much reduced in *H99 EGFR btl/FGFR* mutants, leaving only residual expression in a subset of the invaginated cells (Figure 3C). Thus, EGFR signaling is the predominant factor promoting maintenance of *trh* expression. In its absence, this function can be compensated by Btl/FGFR signaling, as it does during invagination and branching (Figure 3D).

**Figure 3.**
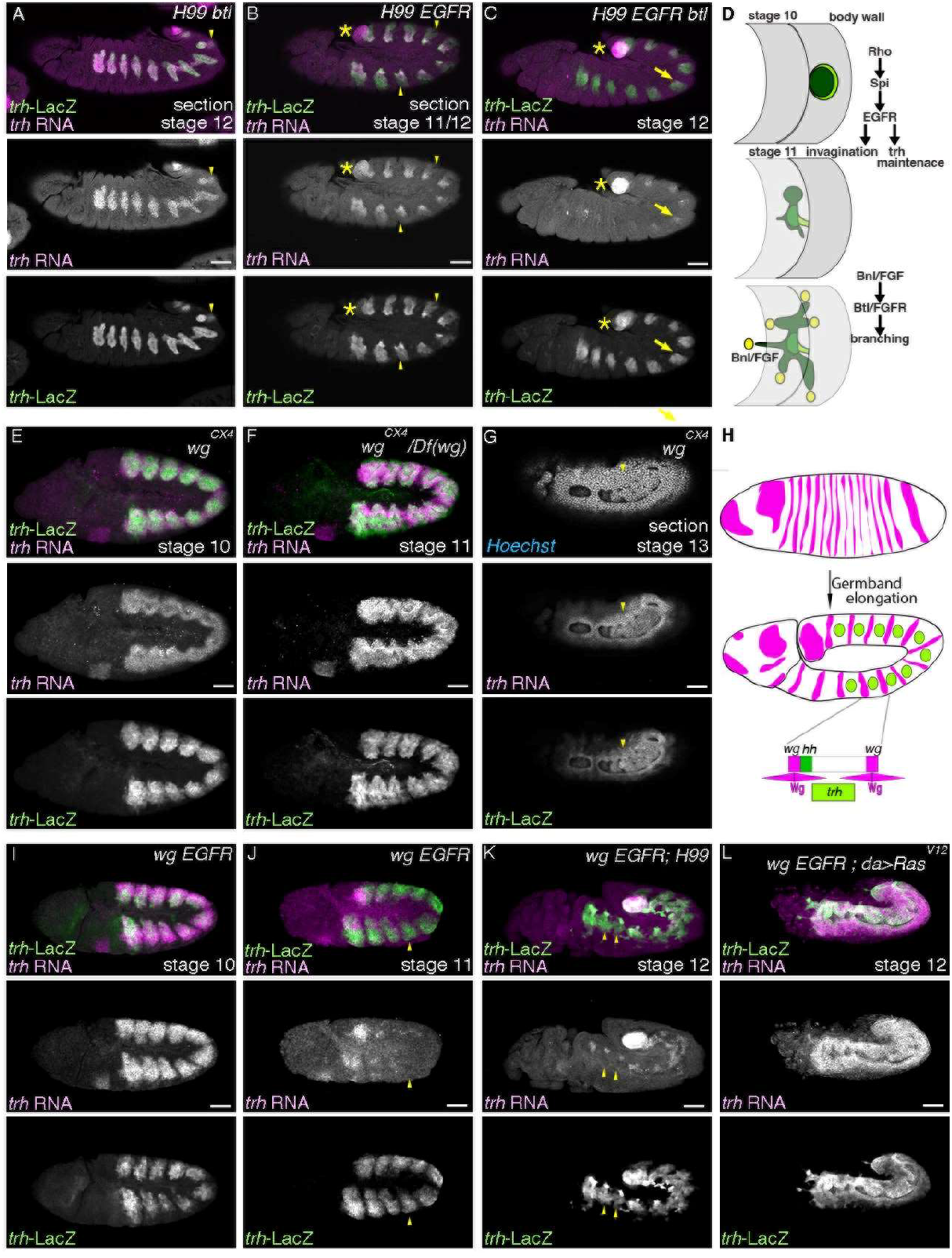
Maintenance of *trh* expression does not follow changes in tissue architecture. Lateral views of embryos stained with *trh*-LacZ and *trh* transcripts. (A-D) In *H99 btl/FGFR* mutants (*H99 btl^Δoh10^ /H99 btl^Δoh24-1^*) (A), *trh* expression is detected both in the distal and the proximal regions whereas in *H99 EGFR* mutants (*top^f2^ /top^f24^; H99*) (B), *trh* expression in the proximal regions is significantly reduced (yellow arrowheads). In *H99 EGFR btl/FGFR* mutants (*top^f2^ /top^f24^; H99 btl^Δoh10^/H99 btl^Δoh24-1^*) (C), residual *trh* expression is detected in parts of the invaginated cells (yellow arrows). Asterisks mark *trh* expression in the posterior spiracle primordia. A schema in D shows stages and main functions of RTK activation in the airway progenitors. (E-L) Phenotypes of *wg/WNT* mutants (E-G) and *wg/WNT EGFR* double mutants (I-L). Wg/WNT is expressed in segmental stripes along the A-P axis (H). Wg/WNT maintains *hh* expression in the posterior neighbors but represses *trh* expression to generate 10 segmentally repeated airway primordia. In the absence of Wg/WNT, the airway progenitor areas expand along the A-P axis (E-G). *trh*-LacZ and *trh* RNA are largely co-expressed since before invagination. Arrowheads in G show that *trh* is expressed in cells that take the 2D planar configuration. In *EGFR wg* double mutants (*top^f24^ wg^CX4^* homozygotes) (I, J), maintenance of *trh* RNA becomes defective at around the stage of invagination (compare I and J). *trh* maintenance is not restored by suppression of apoptosis (*top^f24^ wg^CX4^; H99*) (K) but by *daughterless (da)*-Gal4 driven overexpression of Ras^V12^ (*top^f24^ wg^CX4^; da-Gal4/UAS-Ras^V12^*) (L). Note that *trh* RNA is not detected in cells positive for *trh*-LacZ in J and K (yellow arrowheads). Scale bar is 50 μm

In contrast to a previous model, where maintenance of *trh* expression correlates with the transition of primordial cells from a planar 2D into a 3D tubular tissue architecture (Kondo and Hayashi 2019), we showed that *trh* expression is maintained in the cells on the 2D ectodermal plane in the *hh vvl* double mutants (Figure 1B, Figure 1-figure supplement 1E). Additionally, *trh* expression was very much reduced in the invaginated cells of *H99 EGFR btl/FGFR* mutants (Figure 3C), uncoupling *trh* gene expression from the morphogenetic process of invagination.

To further test if tissue architecture contributes to the maintenance of *trh* expression, we made use of mutations in *wingless (wg)/WNT*, which is expressed in segmental stripes and represses expression of *trh* and *vvl* along the anterior-posterior (A-P) axis (Figure 3E-H) (de Celis et al. 1995; Wilk et al. 1996). In *wg* mutants, some of the progenitors fail to internalize so that cells expressing either distal or proximal markers are detected at the 2D planar embryo surface (Oda et al. 1994; Matsuda et al. 2015b). Correspondingly, *trh* RNA and *trh-*LacZ are detected on the embryo surface in *wg* mutants at stage 13 and even later (Figure 3G).

Similar to *wg* mutants, the initial *trh* expression expands along the A-P axis in *wg EGFR* double mutants (Figure 3I). However, *trh* RNA expression becomes very weak when most cells still reside on the 2D planes around the stage of primordia invagination (Figure 3J). This significant reduction of *trh* expression domain is still evident even in *H99 wg EGFR* mutants where *H99* deficiency suppresses apoptosis (Figure 3K). Together, we conclude that tissue architecture is dispensable for *trh* maintenance.

We suggest instead that *trh* maintenance is stimulated concurrently with the robust EGFR activation that occurs and propagates in the 2D airway primordia (Gabay et al. 1997a; Gabay et al. 1997b; Wappner et al. 1997; Matsuda et al. 2015b). EGFR activation also distalizes gene expression (Matsuda et al. 2015b) and initiates morphogenesis (Brodu and Casanova 2006; Nishimura et al. 2007). In this scenario, rather than indirectly through tubulogenesis (Kondo and Hayashi 2019), RTK signaling directly promotes *trh* maintenance irrespective of tissue geometry. We further note that in double mutants of *aos* and *Gap1*, which are both negative regulators of EGFR signaling, there is expression of airway distal marker on the embryo surface (Matsuda et al. 2015b). This is accompanied by detection of *trh* expression on the embryo surface (Figure 3-figure supplement 1A-D).

Thus, after the Dpp/BMP mediated specification of the airway progenitors along the D-V axis, the radially graded EGFR signaling activation sustains *trh* expression in the progenitors and initiates airway cell differentiation along the P-D axis.

### DRaf/MAPKKK and Dsor1/MAPKK are required for EGFR-mediated priming of the airway progenitors

Trh activity can be boosted by the PI3kinase-PKB pathway through phosphorylation of Serine 665 (Jin et al. 2001). However, it is not clear how far this pathway is required *in vivo* for Trh activity and *trh* autoregulation. We thus investigated if EGFR mediated priming and *trh* expression maintenance in the airway progenitors require the Ras-MAPK pathway (Figure 4A) (Hou et al. 1995; Mishra et al. 2005). When MAPKK-kinase/Draf or MAPK-kinase/Dsor1 is depleted from the embryos with the *ovo-FRT* germline clone technique (Hou et al. 1995; Chou and Perrimon 1996), *trh* expression initiates, as detected with *trh*-LacZ enhancer trap (Figure 4B-C). However, around the stage of primordial invagination, *trh* expression is very much reduced, in both non-invaginated and invaginated cells (Figure 4B-C).

**Figure 4.**
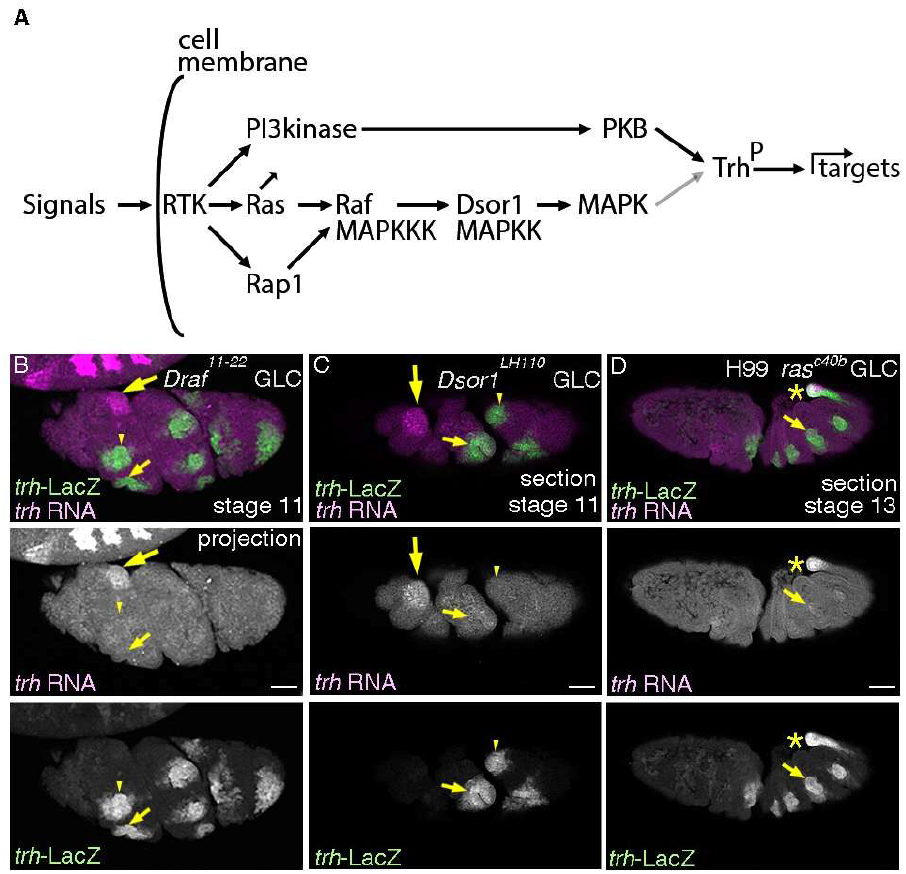
RTKs require the Ras-Raf/MAPKKK-Dsor1/MAPKK pathway for maintenance of *trh* expression. (A) A schema showing that RTK activation involves several downstream signaling branches including the Ras-MAPK pathway and the PI3kinase-PKB pathway. PKB is known to phosphorylate Trh and to upregulate Trh transcriptional activity (black arrow). MAPK may act the same (gray arrow). (B-D) Lateral projection (B) or section (C, D) views of embryos stained with *trh*-LacZ and *trh* transcripts. In the complete absence of Draf/MAPKKK (B) or Dsor1/MAPKK (C), *trh* RNA is hardly maintained in the cells positive for *trh*-LacZ, either invaginated (small yellow arrows) or non-invaginated (yellow arrowheads). In the complete absence of *Drosophila Ras1/Ras85D*, introducing *H99* deficiency suppressing apoptosis allows visualization of airway development till later stages (D), demonstrating residual *trh* expression in parts of the invaginated cells. Large yellow arrows in B and C show *trh* expression in the salivary gland. Yellow asterisks in D mark *trh* expression in the posterior spiracle. Scale bar is 50 μm

### Ras^V12^ boosts Trh activity

*Ras* (*Ras85D* or *Ras1* in *Drosophila*) is a major signaling mediator of RTKs through Draf/MAPKKK-Dsor1/MAPKK-MAPK (Hou et al. 1995; Mishra et al. 2005) and through Pi3kinase-PKB (Figure 4A) (Orme et al. 2006). Consistent with the key role of *Ras85D/Ras1* in RTK signaling, we detected comparable defects in the maintenance of *trh* expression in *EGFR btl/FGFR* double mutants and upon complete loss of *Ras85D/Ras1*, when anti-apoptotic functions are compensated in *H99* chromosomal deletion background (Figure 3C, 4D). Conversely, *trh* maintenance in *EGFR wg/WNT* double mutants upon overexpression of the gain of function form of Ras85D, Ras^V12^ is significantly rescued resembling *wg/WNT* single mutants alone (Fortini et al. 1992) (Figure 3L). Thus, *Ras* serves as the major signaling transducer to promote *trh* maintenance downstream of EGFR and Btl/FGFR.

The mechanisms by which, *Ras* promotes *trh* expression may involve enhancing Trh activity because *trh* auto-regulates its own expression (Figure 4A) (Wilk et al. 1996; Sotillos et al. 2010; Chung et al. 2011). Boosting of Trh activity may involve PKB mediated phosphorylation of Trh (Jin et al. 2001). Alternatively, inferred from the positive roles of MAPK mediated phosphorylation of Hif1, a *trh* homologue (Mylonis et al. 2006), MAPK may directly upregulate Trh activity. Consistently, overexpression of the phosphomimetic form of Trh, Trh^S665D^ or simultaneous double overexpression of Ras^V12^ and Trh^WT^ significantly induced ectopic expression of *trh*-LacZ, whereas single overexpression of Ras^V12^ or Trh^WT^ did not (Figure 5C). The Ras^V12^ induced increase of Trh activity is also detected on the expression of both the distal (*btl* and Gasp/2A12) and the proximal markers (*upd* and *P0144*-LacZ) (Figure 5A-F). We conclude that Ras activation downstream of RTKs is sufficient to upregulate Trh activity toward its downstream targets (Figure 4A).

**Figure 5.**
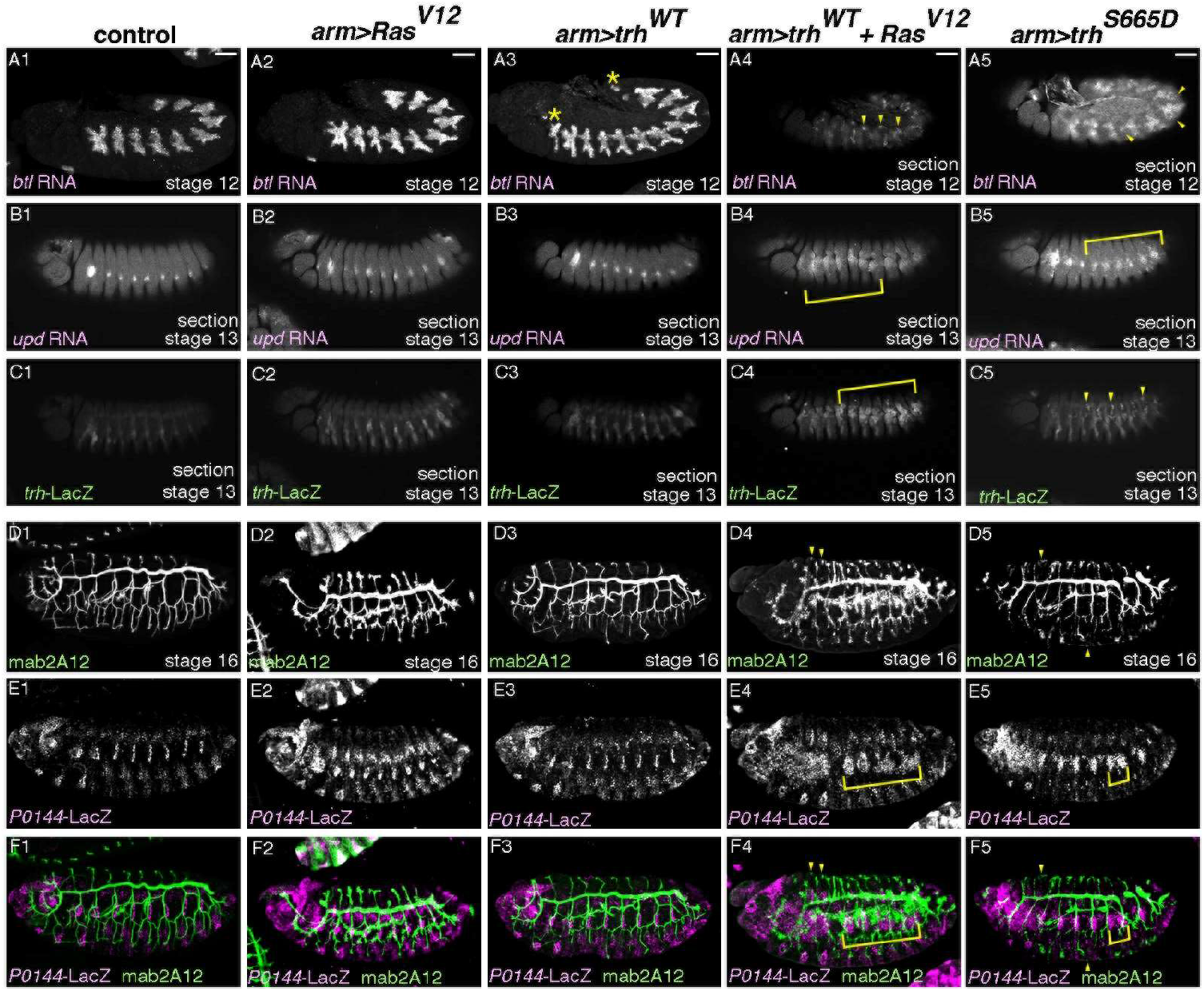
Ras^V12^ boosts Trh activity on downstream genes. (A-F) Lateral views of embryos stained as indicated. Compared to the control (first column), single overexpression of Ras^V12^ (second column) or Trh^WT^ (third column), simultaneous overexpression of Ras^V12^ and Trh^WT^ (fourth column) or single overexpression of Trh^S665D^ (fifth column) induces ectopic expression of *btl* (A), *upd* (B), *trh*-LacZ (C), mab2A12 (Gasp) (D, F) or *P0144*-LacZ (E, F), which are marked by yellow arrowhead or yellow brackets. Asterisks in A3 marks ectopic progenitors in the anterior and the posterior segments. Scale bar is 50 μm.

### Multiple redundant regulators cooperate during early Drosophila airway tubulogenesis

Tubes are common structures in multicellular organisms and tubular organs fulfill essential functions for life (Romero et al. 2025). An active field in stem cell biology engages in artificial organ generation *in vitro* to complement tubular organ dysfunction (Lancaster and Knoblich 2014). In *vivo* studies of tubulogenesis may aid such *in vitro* organoid development. In line with the revised view that a simple master regulator is insufficient for all the steps of the *Drosophila* airway differentiation (de Celis et al. 1995; Llimargas and Casanova 1997; Chen et al. 1998; Boube et al. 2000; Matsuda et al. 2015b; Ogura et al. 2018; Kondo and Hayashi 2019), we characterize the cooperation of both extrinsic and intrinsic, partially redundant regulators to ensure robust airway morphogenesis.

The airway progenitor field specification and its subsequent differentiation is associated with continuous expression of *trh* (Isaac and Andrew 1996; Wilk et al. 1996). *trh* is necessary for expression of all airway differentiation markers and for establishment of the airway tubes (Isaac and Andrew 1996; Wilk et al. 1996; Sotillos et al. 2010; Chung et al. 2011; Matsuda et al. 2015b), assigning it as a master TF of *Drosophila* airway tubulogenesis. Surprisingly, however, *trh* mutant cells invaginate to form tubes, but later revert to a planar configuration (Kondo and Hayashi 2019). Resolving the issue of what promotes progenitor invagination other than Trh, we identified extrinsic Hh and intrinsic Vvl as the primary regulators of invagination, leaving *trh* expression intact.

The airway progenitors are intrinsically defined with at least 3 TFs, *trh*, *vvl* and *grn*. *vvl* defines the central/distal progenitors that invaginate first, whereas *grn* defines the peripheral/proximal progenitors that invaginate later, also reflecting the proximo-distal differences of the airways (Matsuda et al. 2015b). Maintenance of *trh* expression is promoted significantly by autoregulation (Wilk et al. 1996; Sotillos et al. 2010; Chung et al. 2011). However, control of *trh* expression is far more complex, requiring discrete regulations along the A-P, the D-V and the radial/proximo-distal (P-D) axis (Figure 6). Along the D-V axis, our results show that intermediate Dpp/BMP activity promotes maintenance of *trh* expression, whereas high level Dpp/BMP activity is known to be repressive on initiation of *trh* expression (Isaac and Andrew 1996; Wilk et al. 1996). Therefore, different Dpp/BMP levels have two opposing effects on *trh* expression.

**Figure 6.**
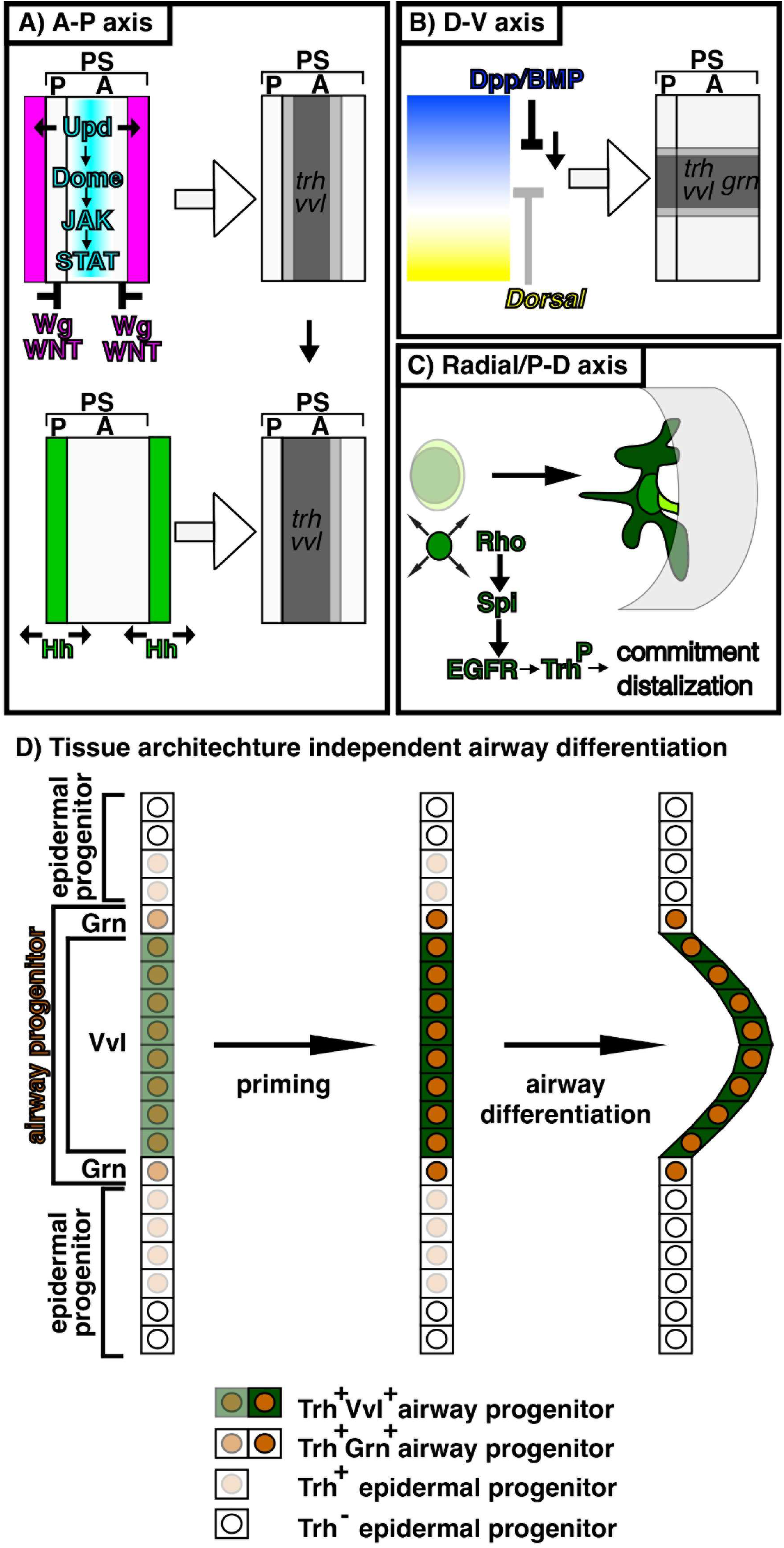
A model for specification and priming of the airway progenitors along the three body axes. Inputs along the A-P axis and the D-V axis generate radial patterning of the airway progenitors to realize the proximo-distal differences of gene expression and morphology. (A) A-P axis; Segments and para-segments (PS) are units of *Drosophila* ectoderm segmentation along the A-P axis (Martinez-Arias and Lawrence 1985). One PS is composed of the posterior of a segment and the anterior of its posterior neighboring segment. Wg/WNT (coloured pink) represses *trh* and *vvl* whereas Upd ligands signal through Domeless-JAK-STAT (coloured blue) to induce them. These activities together generate a graded airway progenitor field along the A-P axis (coloured gray). The *upd* expression domain is a guess. *upd* expression is dynamic (Harrison et al. 1998; Sotillos et al. 2010) and it is not known which *upd* expression is required for inducing the airway primordia. *trh* expression initiates already at stage 8 (Isaac and Andrew 1996) whereas *vvl* expression starts at stage 10. Hh (coloured green) positively regulates expression of *vvl*, thereby distalizing the progenitor field. (B) D-V axis; The ventrally active TF Dorsal (coloured yellow) restricts *dpp/BMP* expression to the dorsal domain. Dpp/BMP (coloured blue) in turn orchestrates the dorsal parts of the embryos. Intermediate level of Dpp/BMP activity promotes expression of *trh*, *vvl* and *grn* to generate a graded airway progenitor field along the D-V axis. (C) P-D axis; The radial patterning is realized by Rho-mediated activation of EGFR. EGFR activation primes the airway progenitors to commit to the airway differentiation program. At the same time, EGFR distalizes the airway differentiation along both gene expression and morphology to establish the P-D axis of the invaginated tubes. Spreading of *rho* expression and promotion of *trh* maintenance may involve EGFR mediated activation of TFs like Trh and Vvl, which boosts their transcriptional activities toward downstream genes including *rho* and *trh*. (D) A model of tissue-architecture-independent regulation of *trh* expression. *trh* expressing cells (brown coloured nucleus) are composed of the epidermal progenitors (pale brown) and the airway progenitors (brown), the latter of which is further classified into *vvl* expressing central/distal cells (green cytoplasm) and *grn* expressing peripheral/proximal cells. Each progenitor type is specified on the 2D cell fields based on the cues along the A-P and the D-V axes (left). Radial EGFR signaling primes the airway progenitors (center, dark brown), to realize morphogenetic and transcriptional differentiation along the P-D axis (right). *trh* expression in the epidermal progenitors ceases based on the epidermal differentiation programs. Airway progenitor invagination may be aided by segregation of the 2 different cell types whereas invagination processes may finetune the cell type determination processes. Note that *grn* expression in the epidermal progenitors and in the early distal progenitors is omitted.

Along the A-P axis, segmentally repeated expression of Unpaired ligands and the resultant activation of Domeless-JAK-STAT signaling precedes and may set, where expression of *trh* and *vvl* initiates (Brown et al. 2001; Sotillos et al. 2010). The airway field is repressed in the embryonic head and the tail by a zinc finger TF, *spalt* (Boube et al. 2000), whereas it is segmentally repressed in the trunk by Wg/WNT (de Celis et al. 1995; Wilk et al. 1996). It is not known what initiates the remaining expression of *trh* and *vvl* in the absence of Unpaired family ligands-Domeless-JAK-STAT signaling (Brown et al. 2001).

Along the radial/P-D axis, the centro-peripheral gradient of Rho-mediated EGFR activation in the 2D progenitor fields promotes maintenance of *trh* expression. Co-expression of Trh and Ras^V12^ potentiates ectopic Trh activity toward downstream genes. Therefore, we suggest that EGFR mediated boosting of TFs like Trh or Vvl may underly *trh* maintenance (Wilk et al. 1996) and the sequential spreading of *rho* expression (Matsuda et al. 2015b), generating a wave of feed-forward loops for progenitor priming and airway differentiation. Additionally, the previously reported *grn*-mediated *trh* maintenance in the peripheral/proximal cells and the *vvl*-mediated enhancement of *trh* expression in the central/distal cells (Matsuda et al. 2015b) argue for region-specific requirements in *trh* maintenance in the primordium.

Our revised model poses that it is pre-determined before invagination, which cells in the *trh* expressing 2D field contribute to airways or to the epidermis (Figure 6D). Only cells receiving appropriate amounts of positive inputs like the intermediate level of Dpp/BMP or sufficient EGFR activation maintain *trh* expression whereas those receiving higher negative inputs like Wg/WNT cease *trh* transcription. The timing of transcriptional silencing and degradation may determine when cells become devoid of *trh* products. Recently, it was discovered that micro-peptides encoded by *polished rice/tarsal-less* promote *trh* expression by suppressing the repressor activity of a zinc finger TF Ovo/Shaven-baby (Mizuno et al. 2026). It is intriguing, where this circuit impinges into the regulatory modes of *trh* expression we described above.

Multiple partially redundant systems on initiation and maintenance of gene expression and morphology sustain development of even simple organs like the *Drosophila* airways. The crucial roles of tubular organs for survival could be the driving force in evolving multiple overlying regulatory schemes, which may secure development of essential organs even when one regulatory scheme becomes dysfunctional.

## Materials and methods

Fly genetics and histochemistry of embryos were done as previously described (Matsuda et al. 2015a; Matsuda et al. 2015b). Marker genes were typically introduced as a single copy, unless otherwise noted.

Fly strains used in this study are:

*aos^Δ7^* (BDSC 1004)

*arm-Gal4* (BDSC 1560)

*btl-Gal4* (a gift from Dr. S. Hayashi)

*D-lacZ* (a gift from Dr. J. Nambu and Dr. S. Russel)

*Draf^11-22^ FRT101* (a gift from Dr. N. Perrimon)

*Dsor1^LH110^ FRT101* (a gift from Dr. N. Perrimon)

*btl^Δoh10^*(Ohshiro and Saigo 1997)

*btl^Δoh24-1^*(Ohshiro and Saigo 1997)

*da-Gal4* (BDSC 8641)

*dpp^H46^*(BDSC 2061)

*dpp^hr92^*(BDSC 2069)

*Dr/msh-lacZ* (a gift from Dr. A. Nose)

*Df(2L)Exel6017=Df(wg)* (BDSC 7503)

*Df(3L)Exel6109=Df(vvl)* (BDSC 7588)

*Gap1^B2^* (a gift from Dr. N. Perrimon)

*GMR19D03* (BDSC 48846)

*grn-GFP* (BDSC 58483)

*H99=Df(3L)H99* (BDSC 1576)

*hh^13C^* (Hosono et al. 2003)

*hh^AC^* (BDSC 1749)

*hs-FLP* (BDSC)

*iroquois-lacZ* (a gift from Dr. S. Campuzano)

*ovoD1 FRT101; hs-FLP* (BDSC 1813)

*ovoD1 FRT82B* (BDSC 2149)

*P0144-lacZ* (a gift from Dr. W. Janning, Flyview)

*Ras85D^Δc40b^* (a gift from Dr. N. Perrimon and Dr. C. A. Berg)

*top^f2^* (BDSC 2768)

*top^f24^* (a gift from Dr. K. Moses)

*trh-lacZ=1-eve-1* (a gift from Dr. N. Perrimon)

*UAS-GFP.nls* (BDSC 4776)

*UAS-trh^WT^/CyO* (a gift from Dr. A. S. Manoukian)

*UAS-trh^S665D^* (a gift from Dr. A. S. Manoukian)

*UAS-Ras^V12^/CyO* (a gift from Dr. G. M. Rubin)

*UAS-Ras^V12^/TM3* (a gift from Dr. N. Perrimon)

*vvl^utH599^* (a gift from Dr. A. Salzberg)

*wg^CX4^* (BDSC 2980)

Antibodies used for immunohistochemistry are;

Rat anti-Dcad (DSHB, donated by T. Uemura) (Oda et al. 1994)

Rabbit anti-GFP (invitrogen)

Rabbit anti-LacZ (Capel)

Rabbit anti-Trh (this study)

Rabbit anti-Vvl (this study)

Double fluorescent labeling with RNA probes and antibodies was carried out as described (Goto and Hayashi 1997).

DNA clones used for in situ RNA detection are;

*caup* (Drosophila Genomics Resource Center, DGRC)

*trh* (Drosophila Genomics Resource Center, DGRC)

*vvl* (a gift from Dr. J. Casanova)

Confocal images were taken by Bio-Rad (Hercules, CA) MRC1024, Olympus (Japan) Fluoview 1000 or Zeiss (Germany) LSM800. Images were processed by ImageJ and figures were prepared with Adobe Photoshop and Illustrator.

## Supporting information

All with suppl.

## Acknowledgements

We thank the members of the fly community who isolated, characterized or distributed fly strains, antibodies or DNA clones. Especially, we thank Drs. C. A. Berg, S. Campuzano, J. Casanova, S. Hayashi, W. Janning, A. S. Manoukian, K. Moses, J. Nambu, A. Nose, N. Perrimon, G.M. Rubin, S. Russel, A. Salzberg, DGRC and BDSC for directly sharing fly strains and DNA clones. We thank Flybase for the *Drosophila* genomic resources. We thank the MBW technical groups and the Stockholm University Imaging Facility. We thank V. Tsarouhas and C. Molenaar for microscope help. Special thanks to Y. Emori and F. Ui-Tei for help in maintaining fly strains after the retirement of K. Saigo.

This work was funded by the Ministry of Education, Culture, Sport, Science and Technology of Japan to K.S. and the Swedish Research Council, the Swedish Cancer Society and German Research Foundation to C.S.

## Author contribution

RM, conceived the project, designed the experiments, performed experiments, interpreted data, drafted the manuscript with inputs from

CH, writing, figure preparation

CH, performed experiments, figure preparation, writing

KS, funding acquisition, provided experiment and analysis tools

CS, funding acquisition, provided experiment and analysis tools, writing

---

**Figure 1-figure supplement 1.**
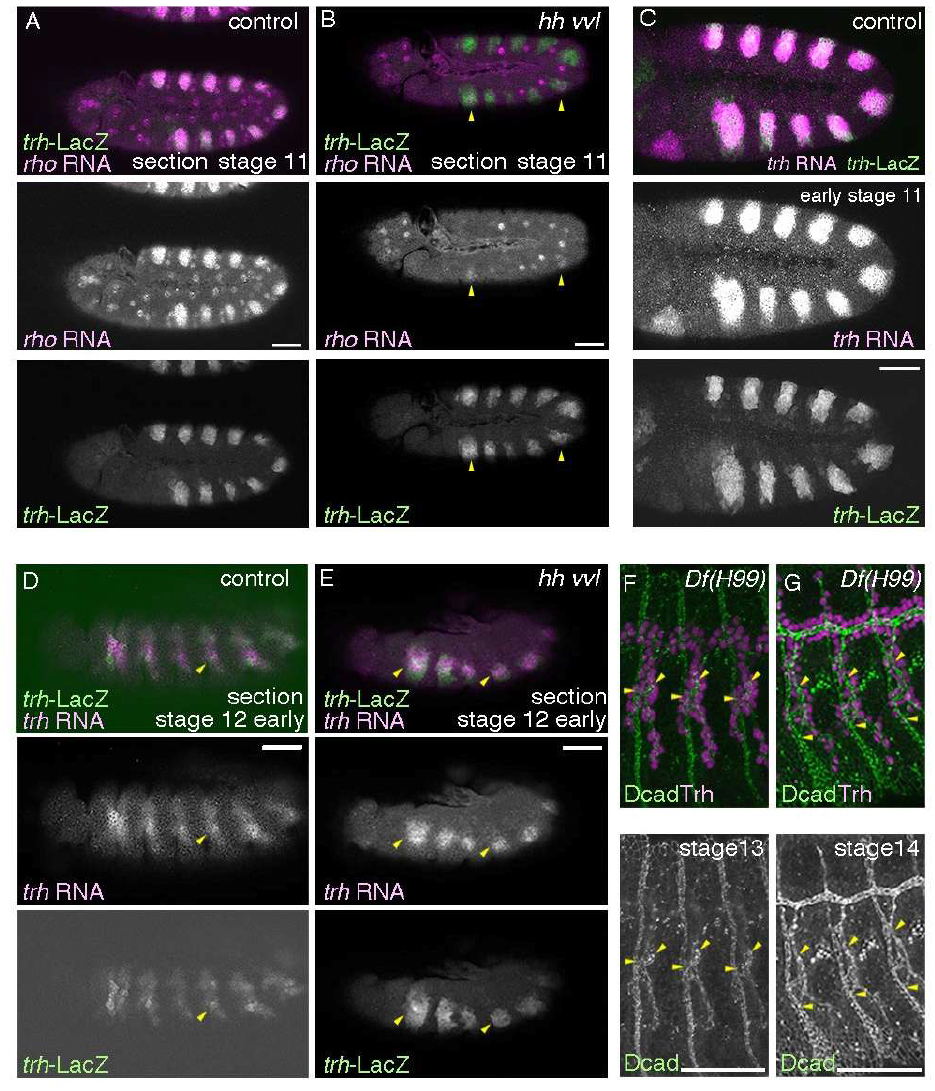
Expression of *rho* and *trh* and the elongation of spiracular branches. (A, B) Compared to the control (A), weak and sporadic expression of *rho* is detected at early stage 11 in *hh vvl* double mutants (*hh^AC^ vvl^utH599^/hh^13C^ Df(vvl)* transheterozygotes) (yellow arrowheads). (C) Expression of *trh* transcripts and *trh*-LacZ largely overlap at late stage 10, which allows tracing the *trh*^+^ cells even after *trh* transcription ceases. (D, E) At early stage 12, *trh* expression is not majorly detected on the embryo surface in the control (D). In contrast, the airway progenitors frequently fail invagination in *hh vvl* double mutants and those cells retain *trh* expression (E). Yellow arrowheads mark the proximal expression of *trh* products. (F, G) Lumens of the spiracular cells significantly elongate between stages 13 and 14, as revealed by double labeling of Trh and Dcad. Yellow arrowheads mark the each edge of the spiracular branches at the epidermis and the TC junctions. Scale bar is 50 μm

**Figure 2-figure supplement 1.**
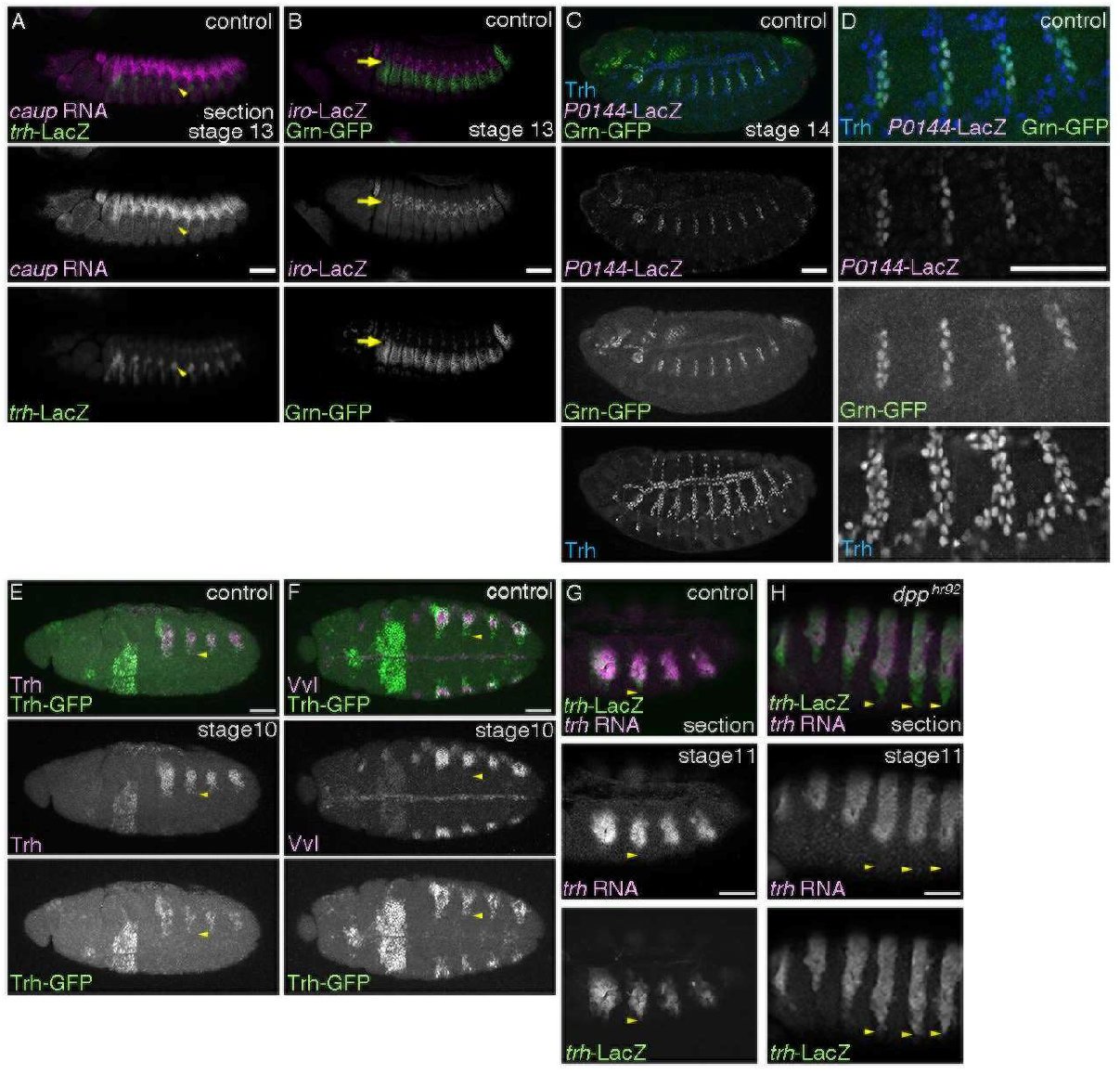
The proximal airways and the ectodermal subdivision. (A-D) Lateral views of stage 13 (A, B) or stage 14 (C, D) embryos. The spiracular branch cells (*trh*-LacZ, yellow arrowheads) are several cells ventrally to the *caup* RNA expressing cells (A). Grn-GFP expresses in the dorso-lateral ectodermal cells that reside ventrally to the *iroquois*-LacZ (*iro*-LacZ)/*araucan*-LacZ) positive dorso-intermediate column (B, yellow arrows). Grn-GFP also expresses in the spiracular branches that are positive for *P0144-*LacZ and Trh (C, magnified in D). (E, F) Ventro-lateral (E) or ventral (F) views of stage 10 embryos stained for Trh-GFP and Trh (E) or Vvl (F). Yellow arrowheads mark the ventral limit of Trh-GFP expression. Note that Trh-GFP expression domains are smaller than those of Trh. (G-H) Lateral views of embryos stained for *trh*-LacZ and *trh* transcripts. At stage 11, *trh*-LacZ positive cells that lose *trh* RNA are pronounced in the ventral parts of the control (G). This becomes more pronounced in the *dpp* hypomorhps (H). Yellow arrowheads mark the ventral limits of *trh*-LacZ expression. A, G and H are single sections. Scale bar is 50 μm

**Figure 3-figure supplement 1.**
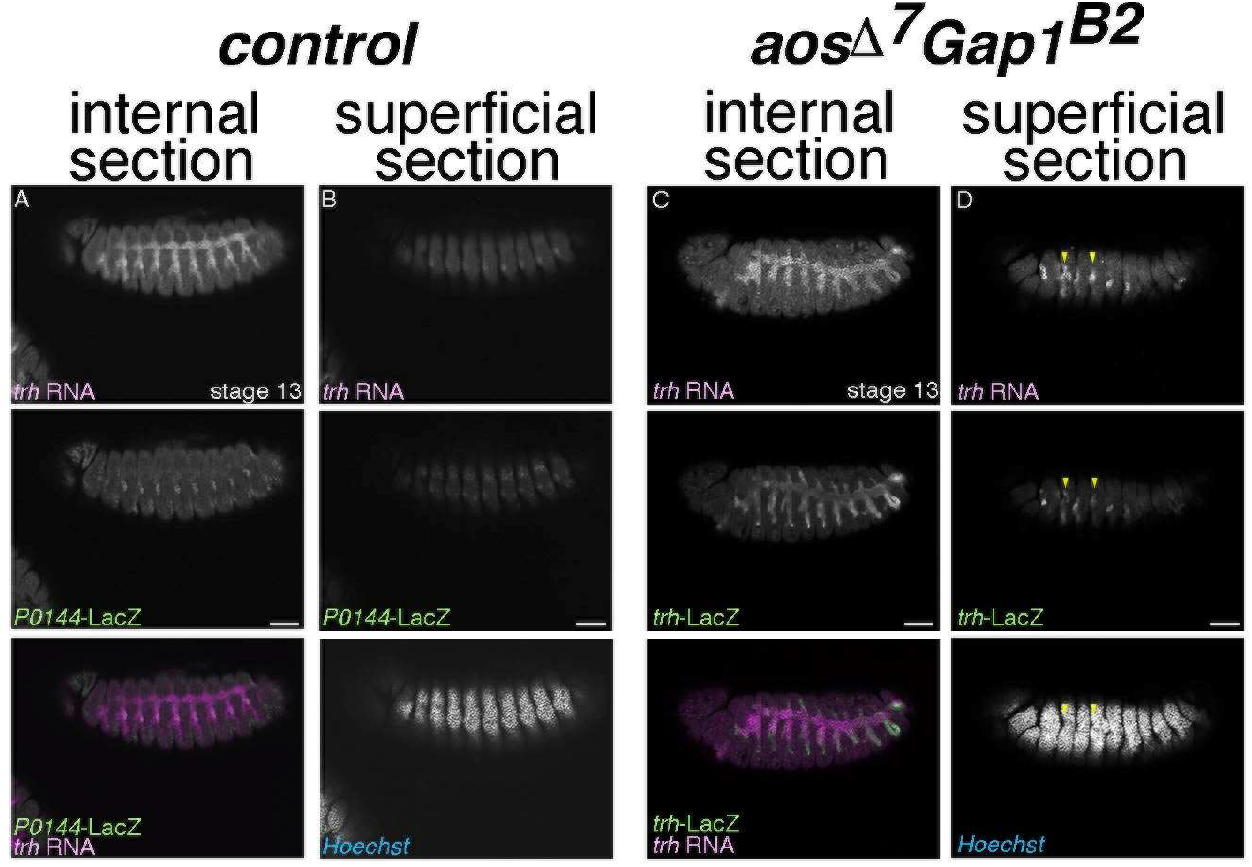
*trh* expression in *aos Gap1* double mutants does not depend on tissue architecture. (A-D) Lateral views of control (A, B) and *aos Gap1* mutant (C, D) embryos stained for *trh* transcripts with *P0144-*LacZ (A, B) or *trh*-LacZ (C, D). Internal (A, C) or superficial sections (B, D). Compared to the control, *trh* expression is often detected in planes at the embryo surface in *aos Gap1* mutant embryos (yellow arrowheads). Scale bar is 50 μm

