## Supplementary material for "Functional definition of the *Drosophila* airway progenitor field through overlapping compensatory regulators": All with suppl.

Fly strains used in this study are:
*aos*$^{\Delta 7}$ (BDSC 1004)

*arm-Gal4* (BDSC 1560)
*btl-Gal4* (a gift from Dr. S. Hayashi)
*D-lacZ* (a gift from Dr. J. Nambu and Dr. S. Russel)
$Draf^{11\text{-}22}$ *FRT101* (a gift from Dr. N. Perrimon)
$Dsor1^{LH110}$ *FRT101* (a gift from Dr. N. Perrimon)
$btl^{\Delta oh10}$ (Ohshiro and Saigo 1997)
$btl^{\Delta oh24\text{-}1}$ (Ohshiro and Saigo 1997)
*da-Gal4* (BDSC 8641)
$dpp^{H46}$ (BDSC 2061)
$dpp^{hr92}$ (BDSC 2069)
*Dr/msh-lacZ* (a gift from Dr. A. Nose)
*Df(2L)Exel6017=Df(wg)* (BDSC 7503)
*Df(3L)Exel6109=Df(vvl)* (BDSC 7588)
$Gap1^{B2}$ (a gift from Dr. N. Perrimon)
*GMR19D03* (BDSC 48846)
*grn-GFP* (BDSC 58483)
*H99=Df(3L)H99* (BDSC 1576)
$hh^{13C}$ (Hosono et al. 2003)
$hh^{AC}$ (BDSC 1749)
*hs-FLP* (BDSC)
*iroquois-lacZ* (a gift from Dr. S. Campuzano)
*ovoD1 FRT101; hs-FLP* (BDSC 1813)
*ovoD1 FRT82B* (BDSC 2149)
*P0144-lacZ* (a gift from Dr. W. Janning, Flyview )
$Ras85D^{\Delta c40b}$ (a gift from Dr. N. Perrimon and Dr. C. A. Berg)
$top^{f2}$ (BDSC 2768)
$top^{f24}$ (a gift from Dr. K. Moses)
*trh-lacZ=1-eve-1* (a gift from Dr. N. Perrimon)
*UAS-GFP.nls* (BDSC 4776)
$UAS\text{-}trh^{WT}/CyO$ (a gift from Dr. A. S. Manoukian)
$UAS\text{-}trh^{S665D}$ (a gift from Dr. A. S. Manoukian)
$UAS\text{-}Ras^{V12}/CyO$ (a gift from Dr. G. M. Rubin)
$UAS\text{-}Ras^{V12}/TM3$ (a gift from Dr. N. Perrimon)
$vvl^{utH599}$ (a gift from Dr. A. Salzberg)
$wg^{CX4}$ (BDSC 2980)

#### Figure and Legends

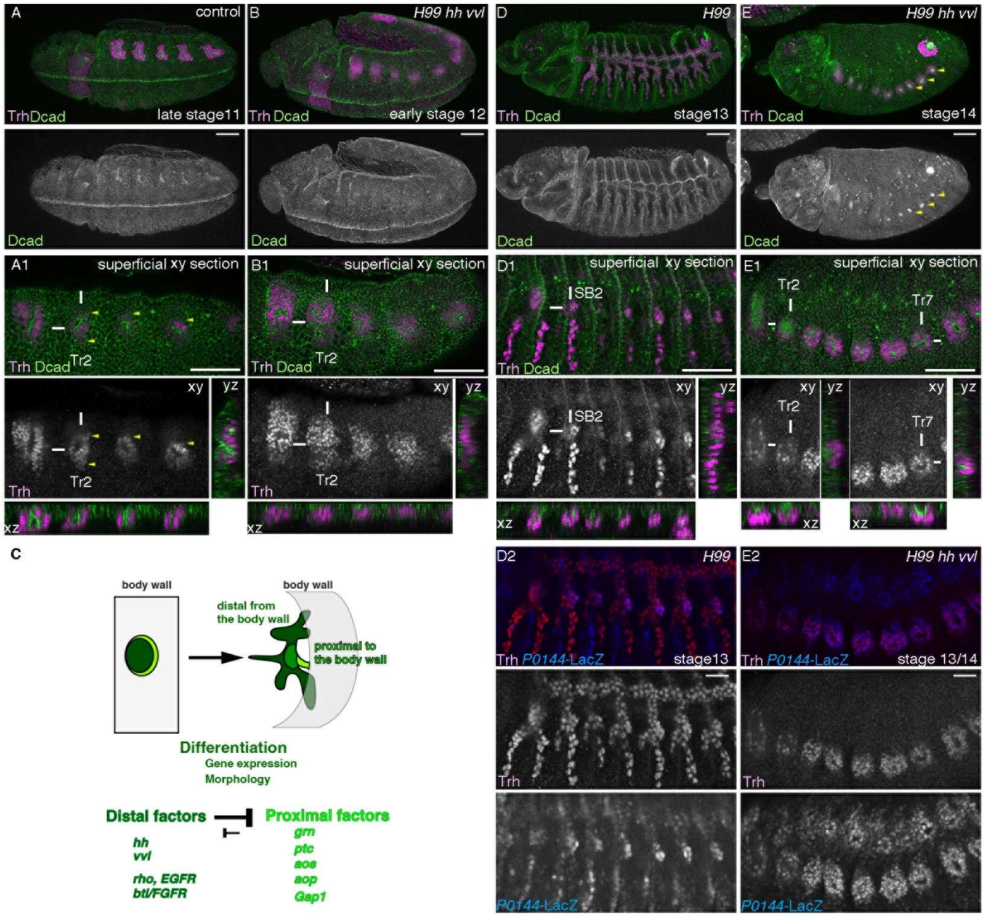


Figure 1

### Figure 1

*hh* and *vvl* empower invagination of the airway progenitors

(A-B, D-E) Progenitor invagination and Trh expression in the control (A, D) or *H99 hh vvl* mutants (*H99* $hh^{13C}$ $vvl^{utH599}$ homozygotes) (B, E), visualized by staining for D-cadherin/Dcad (Shotgun) (Oda et al. 1994; Tepass et al. 1996). A, B, D, D2 and E2 are projections while A1, B1, D1, E1 represent xy- yz- and xz- sections of embryos shown in A, B, D and E, respectively. White lines mark the places of yz- and xz-sectioning. Signals are enhanced in sectional views.

In the control (A, A1), progenitor invagination and tubulogenesis are evident as early as late stage 11. In *H99 hh vvl* mutants at stage 12 (B, B1), apical enrichment of Dcad signals and apical extrusion of $Trh^{+}$ nucleus are irregularly detected. However, persistent progenitor invagination to form deep cavities rarely happens.

Note that in A1, Trh is detected in the epidermal layer (yellow arrowheads) but our reagents do not allow accurate discrimination between epidermal or airway progenitors.

At stages 13/14, compared to the control (D, D1, D2), in *H99 hh vvl* mutants (E, E1, E2), apical enrichment of Dcad signals and apical constriction are persistently detected around the central parts of each primordium, that eventually form shallow cavities (yellow arrowheads in E). Trh and strong *P0144*-LacZ signals overlap mainly in the spiracular branch in the control (D2) whereas in *H99 hh vvl* mutants (E2), *P0144*-LacZ$^+$ progenitors occupy the whole progenitor areas. Trh is detected in smaller areas than *P0144*-LacZ, probably due to the gradual loss of Trh expression in the periphery.
Note that in A1, B1, D1 or E1, the numbers of tracheal metamer (Tr) or spiracular branch (SB) are labelled.
Scale bar is 50 μm

(C) A schema showing that invagination transforms the centro-peripheral patterning of the primordium into the proximo-distal patterning of the tubes. Extrinsic Hh and intrinsic Vvl promote the distal gene expression program (dark green domain) whereas *rho* and *btl/FGFR* activating morphogenesis and distal gene expression programs. The proximal factors including *grn* promote the proximal gene expression program (light green domain).

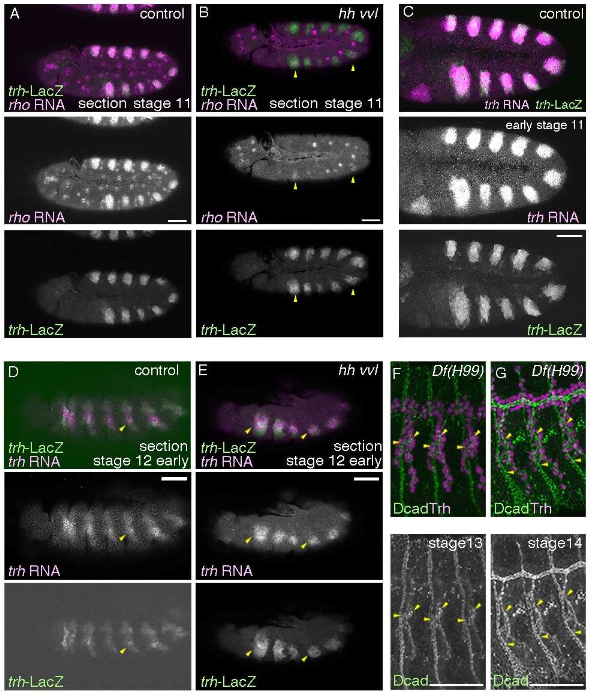

Figure 1-supplement

**Figure 1-figure supplement 1**

Expression of *rho* and *trh* and the elongation of spiracular branches.

(A, B) Compared to the control (A), weak and sporadic expression of *rho* is detected at early stage 11 in *hh vvl* double mutants ($hh^{AC}$ $vvl^{utH599}$/$hh^{13C}$ *Df(vvl)* transheterozygotes) (yellow arrowheads).
(C) Expression of *trh* transcripts and *trh*-LacZ largely overlap at late stage 10, which allows tracing the $trh^{+}$ cells even after *trh* transcription ceases.
(D, E) At early stage 12, *trh* expression is not majorly detected on the embryo surface in the control (D). In contrast, the airway progenitors frequently fail invagination in *hh vvl* double mutants and those cells retain *trh* expression (E). Yellow arrowheads mark the proximal expression of *trh* products.
(F, G) Lumens of the spiracular cells significantly elongate between stages 13 and 14, as revealed by double labeling of Trh and Dcad. Yellow arrowheads mark the each edge of the spiracular branches at the epidermis and the TC junctions.
Scale bar is 50 μm

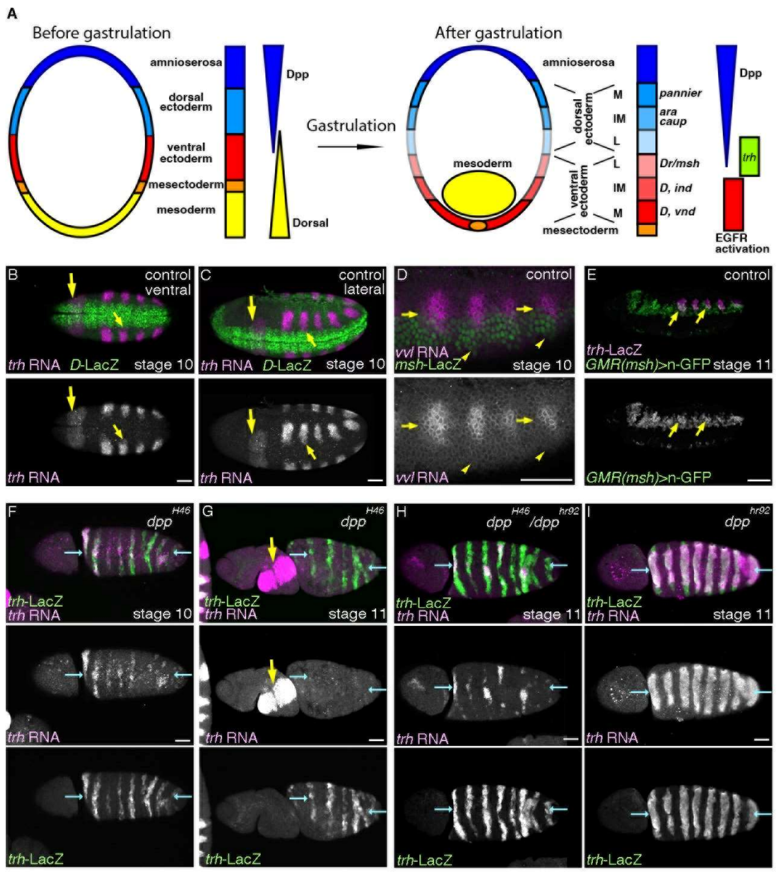


Figure 2

### Figure 2

Intermediate Dpp/BMP activity promotes *trh* expression and the airway progenitors

(A) A schema showing that the embryonic trunk is divided into discrete domains along the D-V axis. Out of the 5 domains (amnioserosa, dorsal ectoderm, ventral ectoderm, mesectoderm and mesoderm), mesoderm invaginates during gastrulation. Graded activities of a TF *dorsal* orchestrates the ventral domains whereas Dpp/BMP signaling directs the dorsal domains. Differential expression of TFs subdivides both the dorsal and the ventral ectoderm into medial (M), intermediate (IM) and lateral (L) columns. EGFR is dynamically activated to establish the ventro-intermediate column (Yagi et al. 1998). Expression of TFs marks the column of dorso-medial (*pannier*), dorso-intermediate (*araucan/ara, caupolican/caup*), ventrol-lateral (*Dr/msh*), ventro-intermediate (*Dicheaete/D, intermediate neuroblasts defective/ind*), ventro-medial (*D, ventral nervous system defective/vnd*).

(B-E) Distribution of *trh* and *vvl* transcripts relative to the ectodermal

subdivision along the D-V axis.
(B-C) Ventral and lateral views of stage 10 embryos. The ventral limit of *trh* expression abuts the dorsal limit of *D*-LacZ expression (small yellow arrows) which marks the ventro-medial and the ventro-intermediate columns. Large arrows show *trh* expression in the salivary gland primordia.
(D-E) *vvl* expression at stage 10 straddles the border between the dorsal ectoderm and the *Dr/msh*-LacZ positive ventro-lateral ectoderm (D, yellow arrows) whereas the ventral limit of *vvl* expression is some cells away from the ventral limit of *Dr/msh*-LacZ expression (D, yellow arrowheads). An enhancer fragment of *Dr/msh* active in the ventro-lateral column marks the ventral parts of the airway tubes at stage 11 (E, yellow arrows).

(F-I) Expression of *trh*-LacZ and *trh* transcripts in allelic series of *dpp* mutants. Dorsal views of embryos where the presumptive dorsal midline are marked with blue arrows.
In *dpp* null mutants (F, G), *trh* expression detected with *trh*-LacZ or *trh* RNA expands to the dorsal midline (blue arrows). However at later stages, *trh* RNA is not maintained though *trh*-LacZ positive cells remain (G). Note that in *dpp* null mutants the body is twisted so that *trh* expression in the salivary gland primordia (yellow arrowheads) and in the airway primordia are simultaneously visible. In the milder condition (H), *trh* RNA is sporadically maintained near the dorsal midline whereas in *dpp* hypomorph homozygotes (I), *trh* RNA is detected in many of the expanded progenitor areas.
Scale bar is 50 µm

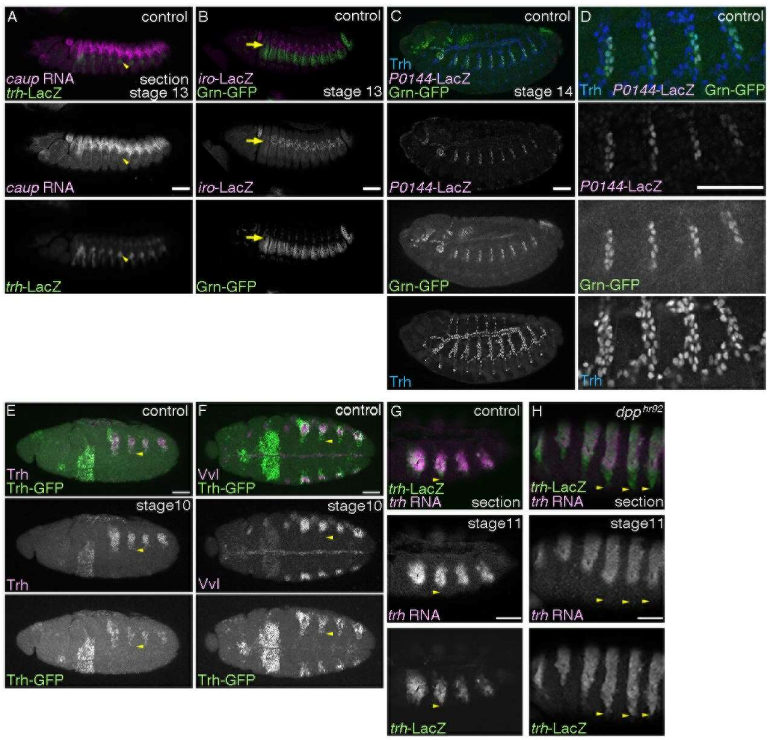


Figure 2-supplement

**Figure 2-figure supplement 1**

The proximal airways and the ectodermal subdivision.

(A-D) Lateral views of stage 13 (A, B) or stage 14 (C, D) embryos. The spiracular branch cells (*trh*-LacZ, yellow arrowheads) are several cells ventrally to the *caup* RNA expressing cells (A). Grn-GFP expresses in the dorso-lateral ectodermal cells that reside ventrally to the *iroquois*-LacZ (*iro*-LacZ)/*araucan*-LacZ) positive dorso-intermediate column (B, yellow arrows). Grn-GFP also expresses in the spiracular branches that are positive for *P0144*-LacZ and Trh (C, magnified in D).

(E, F) Ventro-lateral (E) or ventral (F) views of stage 10 embryos stained for Trh-GFP and Trh (E) or Vvl (F). Yellow arrowheads mark the ventral limit of Trh-GFP expression. Note that Trh-GFP expression domains are smaller than those of Trh.

(G-H) Lateral views of embryos stained for *trh*-LacZ and *trh* transcripts. At stage 11, *trh*-LacZ positive cells that lose *trh* RNA are pronounced in the ventral parts of the control (G). This becomes more pronounced in the *dpp* hypomorhps (H). Yellow arrowheads mark the ventral limits of *trh*-LacZ expression.
A, G and H are single sections.

Scale bar is 50 μm

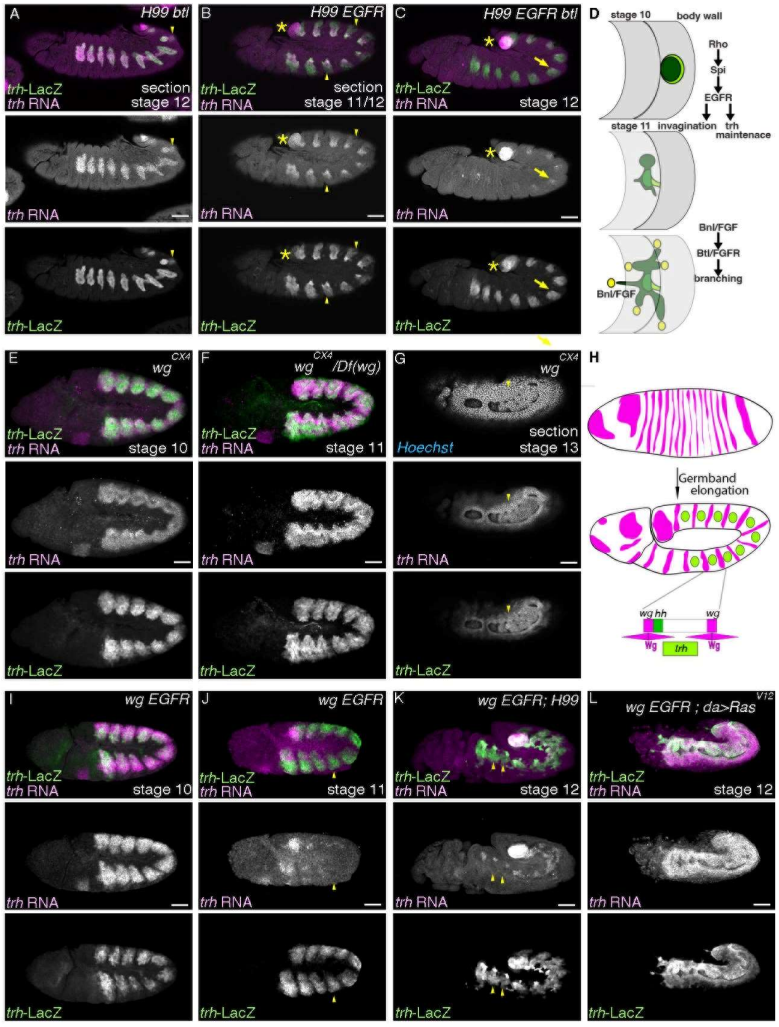


Figure 3

### Figure 3

Maintenance of *trh* expression does not follow changes in tissue architecture

Lateral views of embryos stained with *trh*-LacZ and *trh* transcripts. (A-D) In *H99 btl/FGFR* mutants (*H99* $btl^{\Delta oh10}$ /*H99* $btl^{\Delta oh24\text{-}1}$) (A), *trh* expression is detected both in the distal and the proximal regions whereas in *H99 EGFR* mutants ($top^{f2}$ /$top^{f24}$*; H99*) (B), *trh* expression in the proximal regions is significantly reduced (yellow arrowheads). In *H99 EGFR btl/FGFR* mutants ($top^{f2}$ /$top^{f24}$*; H99* $btl^{\Delta oh10}$/*H99* $btl^{\Delta oh24\text{-}1}$) (C), residual *trh* expression is detected in parts of the invaginated cells (yellow arrows). Asterisks mark *trh* expression in the posterior spiracle primordia. A schema in D shows stages and main functions of RTK activation in the airway progenitors.

(E-L) Phenotypes of *wg/WNT* mutants (E-G) and *wg/WNT EGFR*

double mutants (I-L).
Wg/WNT is expressed in segmental stripes along the A-P axis (H). Wg/WNT maintains *hh* expression in the posterior neighbors but represses *trh* expression to generate 10 segmentally repeated airway primordia. In the absence of Wg/WNT, the airway progenitor areas expand along the A-P axis (E-G). *trh*-LacZ and *trh* RNA are largely co-expressed since before invagination. Arrowheads in G show that *trh* is expressed in cells that take the 2D planar configuration.
In *EGFR wg* double mutants ($top^{f24}$ $wg^{CX4}$ homozygotes) (I, J), maintenance of *trh* RNA becomes defective at around the stage of invagination (compare I and J). *trh* maintenance is not restored by suppression of apoptosis ($top^{f24}$ $wg^{CX4}$ *;H99*) (K) but by *daughterless (da)*-Gal4 driven overexpression of $Ras^{V12}$ ($top^{f24}$ $wg^{CX4}$ *;da-Gal4/UAS-*$Ras^{V12}$) (L). Note that *trh* RNA is not detected in cells positive for *trh*-LacZ in J and K (yellow arrowheads).
Scale bar is 50 μm

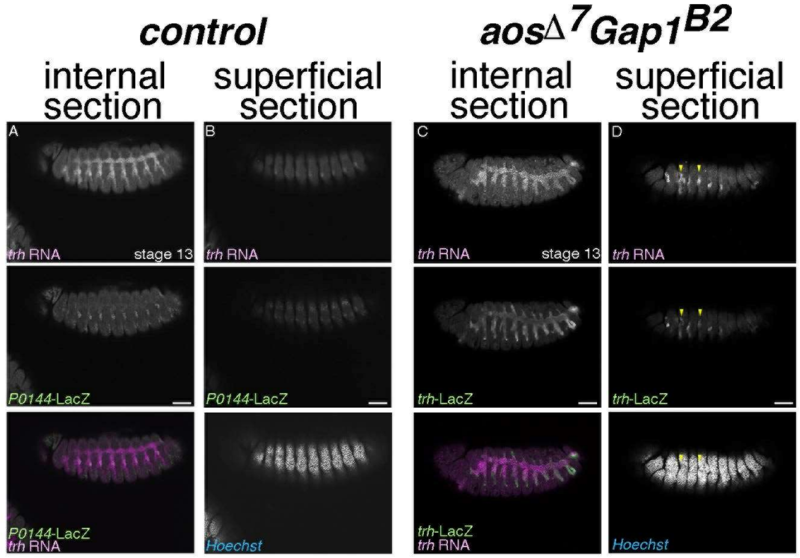

Figure 3-supplement

**Figure 3-figure supplement 1**
*trh* expression in *aos Gap1* double mutants does not depend on tissue architecture.

(A-D) Lateral views of control (A, B) and *aos Gap1* mutant (C, D) embryos stained for *trh* transcripts with *P0144*-LacZ (A, B) or *trh*-LacZ (C, D). Internal (A, C) or superficial sections (B, D). Compared to the control, *trh* expression is often detected in planes at the embryo surface in *aos Gap1* mutant embryos (yellow arrowheads).
Scale bar is 50 µm

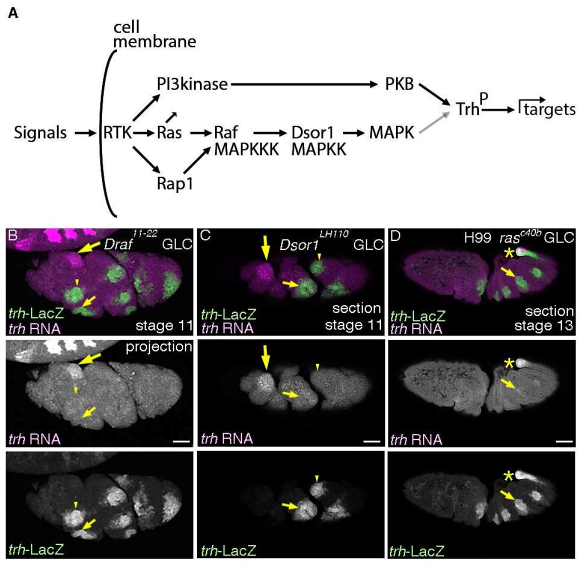


Figure 4

**Figure 4**

RTKs require the Ras-Raf/MAPKKK-Dsor1/MAPKK pathway for maintenance of *trh* expression

(A) A schema showing that RTK activation involves several downstream signaling branches including the Ras-MAPK pathway and the PI3kinase-PKB pathway. PKB is known to phosphorylate Trh and to upregulate Trh transcriptional activity (black arrow). MAPK may act the same (gray arrow).

(B-D) Lateral projection (B) or section (C, D) views of embryos stained with *trh*-LacZ and *trh* transcripts. In the complete absence of Draf/MAPKKK (B) or Dsor1/MAPKK (C), *trh* RNA is hardly maintained in the cells positive for *trh*-LacZ, either invaginated (small yellow arrows) or non-invaginated (yellow arrowheads). In the complete absence of *Drosophila Ras1/Ras85D*, introducing *H99* deficiency suppressing apoptosis allows visualization of airway development till later stages (D), demonstrating residual *trh* expression in parts of the invaginated cells. Large yellow arrows in B and C show *trh* expression in the salivary gland. Yellow asterisks in D mark *trh* expression in the posterior spiracle.
Scale bar is 50 μm

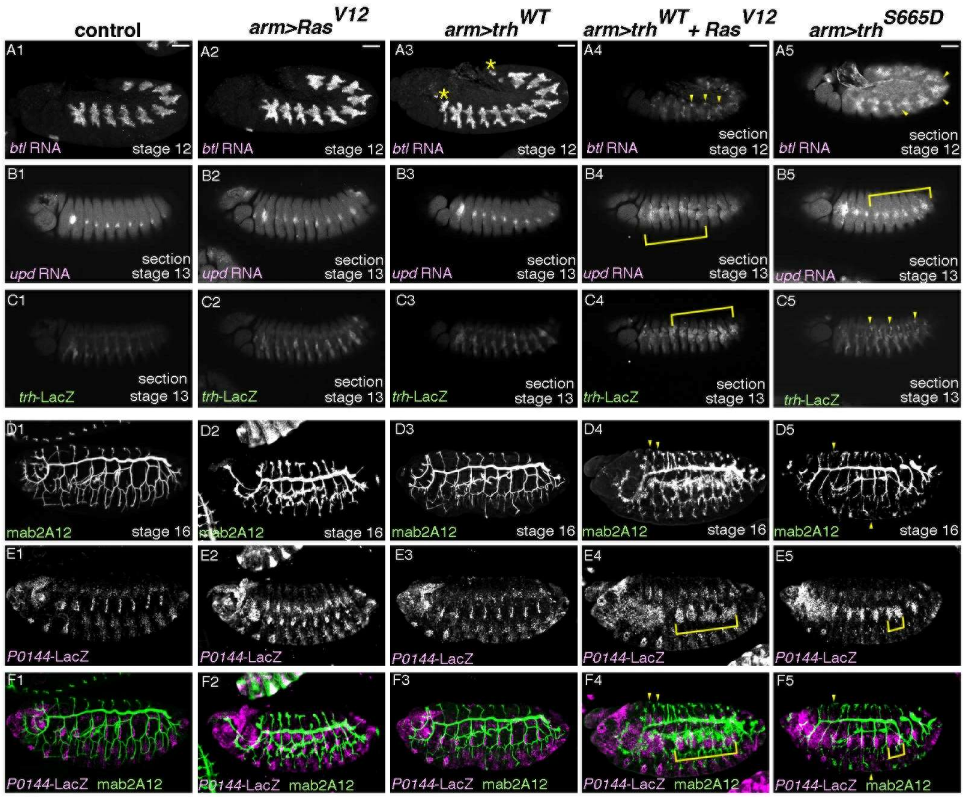


Figure 5

**Figure 5**

Ras$^{V12}$ boosts Trh activity on downstream genes.

(A-F) Lateral views of embryos stained as indicated. Compared to the control (first column), single overexpression of Ras$^{V12}$ (second column) or Trh$^{WT}$ (third column), simultaneous overexpression of Ras$^{V12}$ and Trh$^{WT}$ (fourth column) or single overexpression of Trh$^{S665D}$ (fifth column) induces ectopic expression of *btl* (A), *upd* (B), *trh*-LacZ (C), mab2A12 (Gasp) (D, F) or *P0144*-LacZ (E, F), which are marked by yellow arrowhead or yellow brackets. Asterisks in A3 marks ectopic progenitors in the anterior and the posterior segments.
Scale bar is 50 µm.

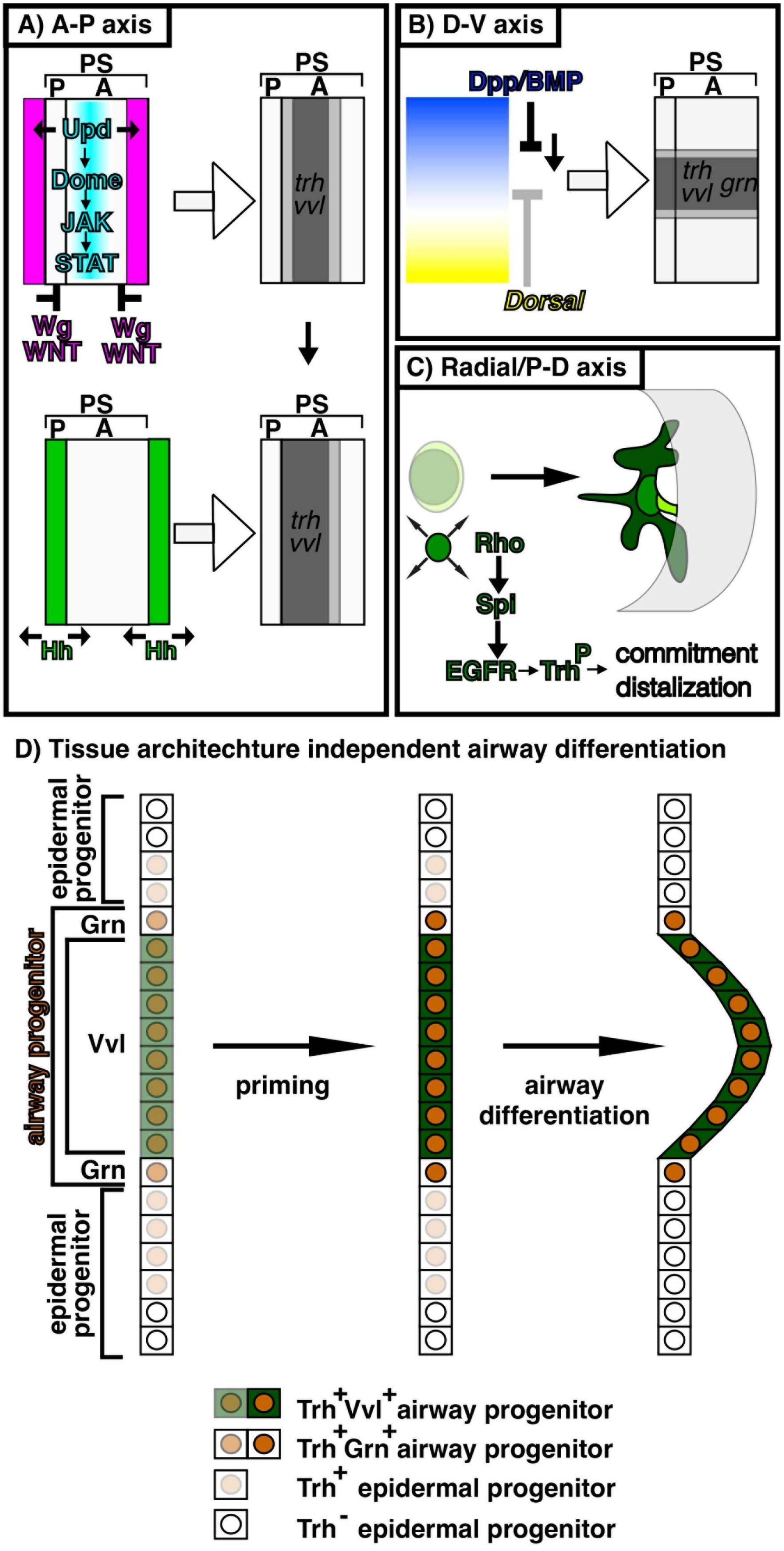
A) A-P axis
PS
P A
Upd
Dome
JAK
STAT
Wg
WNT
Wg
WNT
trh
vvl
Hh
Hh
B) D-V axis
Dpp/BMP
Dorsal
trh
vvl grn
C) Radial/P-D axis
Rho
Spl
EGFR→Trh P → commitment
distalization
D) Tissue architechture independent airway differentiation
epidermal progenitor
airway progenitor
Grn
Vvl
Grn
epidermal progenitor
priming
airway differentiation
Trh+Vvl+airway progenitor
Trh+Grn+airway progenitor
Trh+ epidermal progenitor
Trh- epidermal progenitor

Figure 6

**Figure 6**
A model for specification and priming of the airway progenitors along the three body axes

Inputs along the A-P axis and the D-V axis generate radial patterning of the airway progenitors to realize the proximo-distal differences of gene expression and morphology.
